# Protein structure analysis of the interactions between SARS-CoV-2 spike protein and the human ACE2 receptor: from conformational changes to novel neutralizing antibodies

**DOI:** 10.1101/2020.04.17.046185

**Authors:** Ivan Mercurio, Vincenzo Tragni, Francesco Busco, Anna De Grassi, Ciro Leonardo Pierri

**Author notes:** **Abbreviations:** Severe acute respiratory syndrome, SARS; coronavirus CoV; receptor binding domain, RBD; angiotensin converting enzyme 2, ACE2; fragment antigen binding, FAB; Word Health Organizzation, WHO; complementary-determining regions, (CDR); SwissPDBViewer, SPDBV. These authors equally contributed (Ivan Mercurio, Vincenzo Tragni).

## Abstract

The recent severe acute respiratory syndrome, known as Corona Virus Disease 2019 (COVID-19) has spread so much rapidly and severely to induce World Health Organization (WHO) to declare state of emergency over the new coronavirus SARS-CoV-2 pandemic. While several countries have chosen the almost complete lock-down for slowing down SARS-CoV-2 spread, scientific community is called to respond to the devastating outbreak by identifying new tools for diagnosis and treatment of the dangerous COVID-19. With this aim we performed an *in silico* comparative modeling analysis, which allows to gain new insights about the main conformational changes occurring in the SARS-CoV-2 spike protein, at the level of the receptor binding domain (RBD), along interactions with human cells angiotensin converting enzyme 2 (ACE2) receptor, that favour human cell invasion. Furthermore, our analysis provides i) an ideal pipeline to identify already characterized antibodies that might target SARS-CoV-2 spike RBD, for preventing interactions with the human ACE2, and ii) instructions for building new possible neutralizing antibodies, according to chemical/physical space restraints and complementary determining regions (CDR) mutagenesis of the identified existing antibodies. The proposed antibodies show *in silico* a high affinity for SARS-CoV-2 spike RBD and can be used as reference antibodies also for building new high affinity antibodies against present and future coronavirus able to invade human cells through interactions of their spike proteins with the human ACE2. More in general, our analysis provides indications for the set-up of the right biological molecular context for investigating spike RBD-ACE2 interactions for the development of new vaccines, diagnosis kits and other treatments based on the usage or the targeting of SARS-CoV-2 spike protein.

## INTRODUCTION

Scientific community is called to respond to a pandemic of respiratory disease that has spread with impressive rate among people of all the world. The new coronavirus has been called SARS-CoV-2 and the related disease indicated as COVID-19. WHO reports that positive patients in the world are 1353361 with 79235 (April 9^th^, 2020) ascertained died people due to COVID-19 complications. It also appears that these numbers might be a smaller number of real cases due to our inability in quantifying rescued or asymptomatic people.

In order to limit death rate and SARS-CoV-2 spread, it needs to develop a vaccine and to identify new small molecules able to prevent or treat COVID-19 complications, as well as to prepare new quick diagnosis kits, able to quantify the real number of people exposed to SARS-CoV-2. Among the main actors of SARS-CoV-2 infection the SARS-CoV-2 spike proteins, RNA dependent RNA polymerases and proteases deserve to be mentioned. Indeed, RNA dependent RNA polymerase has become one of the main targets of a nucleoside analog antiviral drug, the remdesivir, already used for reducing complications due to Ebola, Dengue and MERS-CoV infections (1–6). At the same time viral protease inhibitors (7–10) are under investigation for their ability in preventing virus protein (with specific reference to spike proteins) cleavage (11) leading to the fusion of virus proteins with host cell membranes. Also anti-inflammatory antibodies/drugs in combination with anticoagulant molecules are under investigation for limiting coagulopathies (12–15) and cytokine signaling impressively triggered by SARS-CoV-2 infection (16–20). Finally, the same SARS-CoV-2 spike protein has become the most investigated target due to its ability in forming interactions with the human ACE2 receptor, causing fusion events that make possible for the virus to penetrate host human cells (21–23).

The crucial role played by the spike protein is also due to the possibility to use the recombinant SARS-CoV-2 spike protein for triggering immune response, working as a vaccine, that may help in preventing and treating COVID-19, similarly to what proposed recently (24–26).

For clarifying SARS-CoV-2 infection mechanisms, several research groups have recently solved the crystallized structure of the entire SARS-CoV-2 spike protein (6vsb.pdb (21); 6vxx.pdb and 6vyb.pdb (22)), in pre-fusion conformation, and/or SARS-CoV-2 spike RBD domain in complex with the human ACE2 (6vw1.pdb; 6lzg.pdb).

In light of the available cited crystallized/cryo-em solved structures, here we propose a strategy for identifying/drawing new SARS-CoV-2 antibodies directed against the RBD of SARS-CoV-2 that could be used for contrasting SARS-CoV-2 infection, aiming to prevent pre-/post-fusion spike conformation interconversion, responsible for virus invasion, and to provide a molecular structural context for studying new diagnosis kits based on the interactions between our engineered antibodies and the human SARS-CoV-2 spike RBD.

## MATERIALS AND METHODS

### 2.1. Crystal Structure Sampling Via Folding Recognition and multiple sequence alignments (MSA)

CoV-Spike and ACE2 homologus protein-crystallized structures were searched by using the folding recognition methods implemented in pGenThreader and i-Tasser according to our validated protocols (27–30).

The sequences of the retrieved 48 crystallized structures (with reference to those crystallized structures indicated with “Certain” or “High” confidence level in pGenThreader output) were aligned by using ClustalW (31) implemented in the Jalview package (32). The 3D coordinates from the 48 crystallized structures were superimposed for comparative purposes by using PyMOL (33) according to our validated protocols (29, 34).

Protein-protein and protein-ligand binding regions were highlighted by selecting residues within 4 Å at the protein-protein interface or from the investigated ligands, in the superimposed structures. All the generated 3D all atom models were energetically minimized by using the Yasara Minimization server (35).

### 2.2. 3D atomic models preparation of SARS-CoV-2 Spike protein in post-fusion conformation and SARS-CoV-2 Spike-ACE2 interactions in pre-fusion conformations

The 3D comparative model of SARS-CoV-2 spike trimer in post-fusion conformation was built by multi-template modeling by using Modeller (36). More in detail, the human SARS-CoV-2 spike protein sequence was aligned to the sequences of the available entire post fusion conformation of the murine coronavirus spike protein (6b3o, (37)) and the remaining available crystallized subdomains of other coronavirus spike proteins in post fusion conformations (5yl9.pdb (38); 1wyy.pdb (39) and 1wdf (40)). Sequences of the cited crystallized structure fragments were used as query sequences for sampling the corresponding entire spike monomer sequences, by reciprocal-blastp, to be aligned with sequences of the investigated structures for comparative purposes. The obtained MSA was used for driving the multi-template modeling.

Then a complex 3D model representing the pre-fusion spike trimer interacting with three ACE2 functional receptor units was built by superimposing the recently solved cryo-EM prefusion structure of SARS-CoV-2 spike trimer complex (6vsb.pdb, (21); 6vyb.pdb and 6vxx.pdb (22)), the SARS-CoV-2 spike RBD crystallized in complex with the human ACE2 (6vw1.pdb; 6lzg.pdb) the SARS-CoV-1 spike trimer interacting with one ACE2 functional receptor (conformations 1-3, 6acg.pdb, 6acj.pdb, 6ack.pdb, (41) and 6cs2.pdb, (42)), the SARS-CoV-1 spike-RBD crystallized in complex with the human ACE2 (2ajf.pdb, (43)).

For investigating pre-/post-fusion conformation interconversion we superimposed the pre-fusion available crystallized structures of SARS-CoV2 spike proteins and the generated 3D models about pre-fusion conformation of the spike trimer in complex with three ACE2 units, to the obtained 3D model of the post fusion conformation. All the generated 3D all atom models were energetically minimized by using the Yasara Minimization server (35)

### 2.3 Antibody 3D modeling and mutagenesis

Starting from the 3D atomic coordinates of the crystallized neutralizing antibodies m396 (2dd8.pdb(44)) and S230 (6nb7.pdb, (45)) directed against the SARS-CoV-1 spike RBD domain, we modelled the interactions of m396 and S230 (6nb7.pdb, (45)) with SARS-CoV-2 spike RBD domain, by superimposing the fragment antigen based (FAB) portions of m396 (2dd8.pdb (44)) and S230 (6nb7.pdb, (45) (both complexed with SARS-CoV-1 RBD) with the SARS-CoV-2 spike RBD domain, complexed with ACE2 (6vw1.pdb), by using PyMOL.

For creating a more specific antibody directed against SARS-CoV-2 spike RBD, we replaced residues of the CDR regions of the m396 crystallized FAB portion with residues that may complement and fulfill better the SARS-CoV-2 RBD surface. Mutagenesis analyses and modeling of the incomplete residues within the crystallized structures were performed by using SPDBV (46) and/or PyMOL (47).

The proposed complete IgG chimeric antibodies were obtained by superimposing the above cited m396 and the resulting engineered FAB portions, in complex with SARS-CoV-1/2 RBD, to the 3D atomic model of a crystallized IgG, available on the PDB (1igt.pdb, (48)) by using SPDBV and PyMOL, according to our validated protocols (29, 30, 49), that allow to model missing residues, solving clashes and breaks in the backbone.

Each glycosylation ladder coming from the crystal structures here investigated (1igt.pdb; 2dd8.pdb; 6nb7.pdb) was alternatively retained within the generated structural models.

After superimposition operations, allowing backbone connections, we renumbered all the atoms and the residues present in the resulting final pdb file, by using an in-house developed Perl script. The obtained final models were examined in VMD, PyMOL, and SPDBV according to our protocols (30, 49). All the generated 3D all atom models were energetically minimized by using the Yasara Minimization server (35).

### 2.4. Energy calculations

The FoldX AnalyseComplex assay, was performed to determine the interaction energy between the four generated antibodies and the RBD domains of SARS-CoV-1/2 spike proteins, but also for determining the interaction energy between ACE2 and the interacting spike RBDs for comparative purposes.

The way the FoldX AnalyseComplex operates is by unfolding the selected targets and determining the stability of the remaining molecules and then subtracting the sum of the individual energies from the global energy. More negative energies indicate a better binding. Positive energies indicate no binding (50, 51). The energy calculated for the crystallized m396-SARS-CoV-1 RBD protein complex was used as a reference value.

## RESULTS

### 3.1. Modelling of the SARS-CoV-2 spike protein in post-fusion conformation

The main event that allows virus envelop fusion with the host human cell plasma membrane concerns a conformational change occurring at the SARS-CoV-2 spike protein that converts from pre-fusion conformation to post-fusion conformation after interactions with ACE2 and spike protein cleavage. While SARS-CoV-2 spike protein trimer has been resolved by cryo-em (6vsb.pdb (21); 6vxx.pdb and 6vyb.pdb (22)), the post-fusion conformation is not available, yet. According to (11) Coutard et al., protein cleavage at site S1/S2 and S2’ produces the division of the spike protein in two subdomains, i.e. the N-ter S-I ectodomain (containing the RBD interacting with ACE2) and the C-ter S-II membrane anchored subdomain, forming the SARS-CoV-2 spike protein in post fusion conformation, able to trigger the fusion of the viral envelope with host cell plasma membrane determining host cell invasion.

For modelling 3D post-fusion conformation of SARS-CoV-2 spike protein we searched for SARS-CoV-2 spike protein homologous structures and found 48 crystallized structures that included poses of the whole SARS-CoV-2 spike proteins or about protein domains of SARS-CoV-2 spike proteins in complex with protein interactors (i.e. ACE2), several pre-fusion conformations of other coronavirus spike proteins, one coronavirus spike protein in post-fusion conformation and three further protein subdomains about spike proteins in post fusion conformation (Supp. Tab. 1).

Thus, we built a MSA by aligning the sequence of the human SARS-CoV-2 spike protein, the sequence of the available whole post fusion conformation of the murine coronavirus spike protein (6b3o, (37)), sequences of the remaining crystallized subdomains of other coronavirus spike proteins in post fusion conformations (5yl9.pdb (38); 1wyy.pdb (39) and 1wdf (40)), together with their complete counterpart sequences sampled by reciprocal-blastp (Fig. 1).

**Fig. 1.**
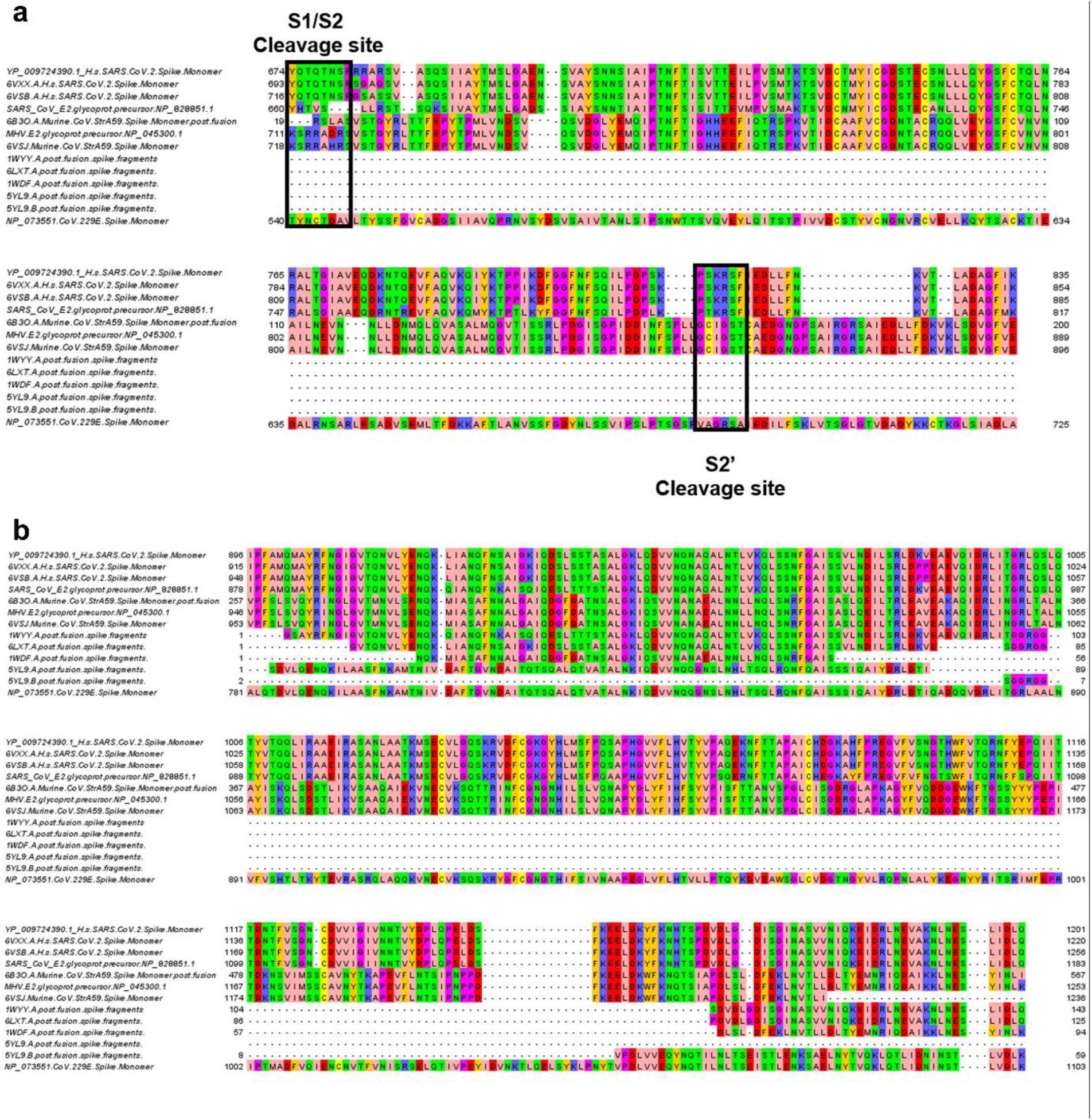
Extract of the MSA of SARS-CoV-2 spike protein monomer with the sequences of crystallized structures of the spike whole protein or protein fragments observed in the post-fusion conformation from other coronavirus, resulting from sequence cleavage. Panel a. Black boxes indicate the position of cleavage sites. In panel “a” it is reported the 6b3o.pdb based sequence-structure alignment used for modeling the first portion of SARS-CoV-2 spike protein in post fusion conformation (amino acids S704-771A, YP_009724390.1 residues numbering). In panel “b” it is reported the 6b3o.pdb based sequence-structure alignment used for modeling the second portion of SARS-CoV-2 spike protein in post fusion conformation (amino acids 922-1147, YP_009724390.1 residues numbering).

In the provided MSA (Fig. 1) it is possible to observe the conserved S1/S2 and S2’ cleavage sites, according to (11) and the sequence of the C-terminal domain resulting from the cleavage. Starting from the cited multi-template sequence alignment and according to our validated protocols about multi-template 3D modeling (30, 36), we built the 3D model of a monomer of SARS-CoV-2 spike protein in post fusion conformation (Fig. 2). The modelled SARS-CoV-2 spike post-fusion conformation consists of residues 704-771 and 922-1147, YP_009724390.1 residues numbering, resulting from protein cleavage (11) and also the only protein fragments with a solved structure in 6b3o.pdb aligned (aminoacids 741-807 and 972-1248, NP_045300.1/6b3o.pdb residues numbering) counterpart (37).

**Fig. 2.**
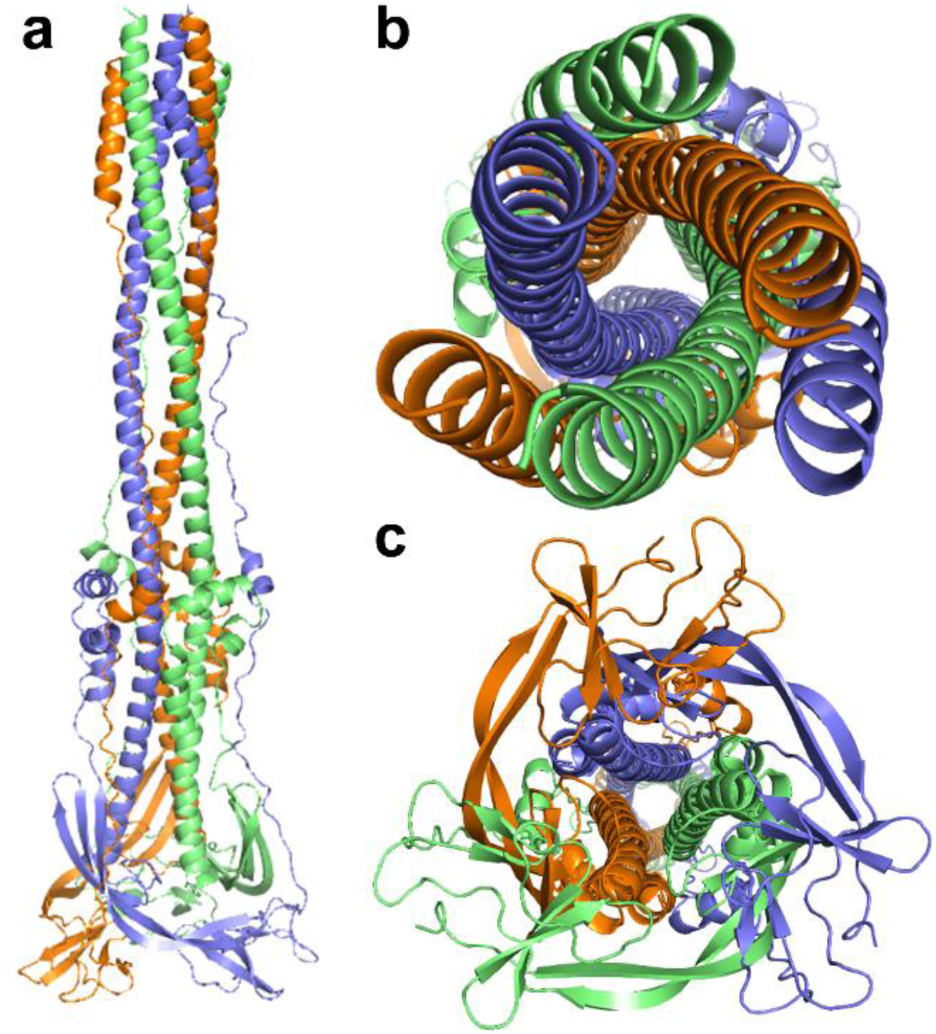
SARS-CoV-2 spike protein (S-II domain) 3D model in post fusion conformation. Lateral view (panel a), top view (panel b) e bottom view (panel c) of the SARS-CoV-2 spike protein trimer 3D comparative model, reported in cartoon colored representation.

The trimer of the SARS-CoV-2 spike protein in post fusion conformation was obtained by duplicating two times the obtained monomer and superimposing the three SARS-CoV-2 spike protein monomers on the three SARS-CoV-1 spike protein monomers reported in 6b3o.pdb (Fig. 2). The 3D comparative model of SARS-CoV-2 spike protein trimer built by multi-template comparative modeling showed an RMSD lower than 0.5 Å with the murine coronavirus spike protein in post-fusion conformation (6b3o.pdb). The resulting model (Fig. 2) appeared elongated and narrow, according to what observed in fragments of the spike proteins crystallized in post-fusion conformations, whose sequences are reported in Fig. 1 and whose PDB_ID are listed in Supp. Tab. 1.

### 3.2. Modelling of the interactions between the SARS-CoV-2 spike protein and the human ACE2 along pre-/post-fusion conformation interconversion

Among the sampled crystallized structures, it was possible to observe three PDB_ID about the entire SARS-CoV-2 spike proteins and two about SARS-CoV-2 spike RBD protein interacting with the human ACE2 (Supp. Tab. 1). Furthermore, it was possible to highlight several crystallized structures about SARS-CoV-1 and MERS-CoV spike proteins as single proteins or in complex with their receptors or dedicated antibodies (Supp. Tab. 1). Notably, among the sampled structures, also the four entries used for building the 3D comparative model of the post-fusion conformation, were sampled (Supp. Tab. 1).

For modeling main interactions occurring between SARS-CoV-2 spike proteins and ACE2, thanks to the high percentage of identical residues shared by spike RBD from several CoV strains (Fig. 3), it was possible to structurally align three objects consisting of the human ACE2-SARS-CoV-1 spike-RBD protein complex (2ajf.pdb) to the human ACE2-SARS-CoV-2 spike-RBD protein complex (6vw1.pdb, 6lzg.pdb) and the SARS-CoV-2 spike protein trimer (6vsb.pdb; 6vxx.pdb; 6vyb.pdb). More in detail, the superimposition performed by using PyMol was leaded by the structural alignment of the RBD of ACE2-SARS-CoV1 (2ajf.pdb) and ACE-2-SARS-CoV-2 (6vw1.pdb, 6lzg.pdb) spike proteins (Fig. 4), followed by the structure alignment with SARS-CoV-2 spike protein trimer (6vsb.pdb; 6vxx.pdb; 6vyb.pdb). Notably, we obtained an efficient superimposition of the two RBD domains (RMSD lower than 0.5 Å) of the human SARS-CoV-1 and SARS-CoV-2 spike proteins also due to their high percentage of identical residues (> 75%).

**Fig. 3.**
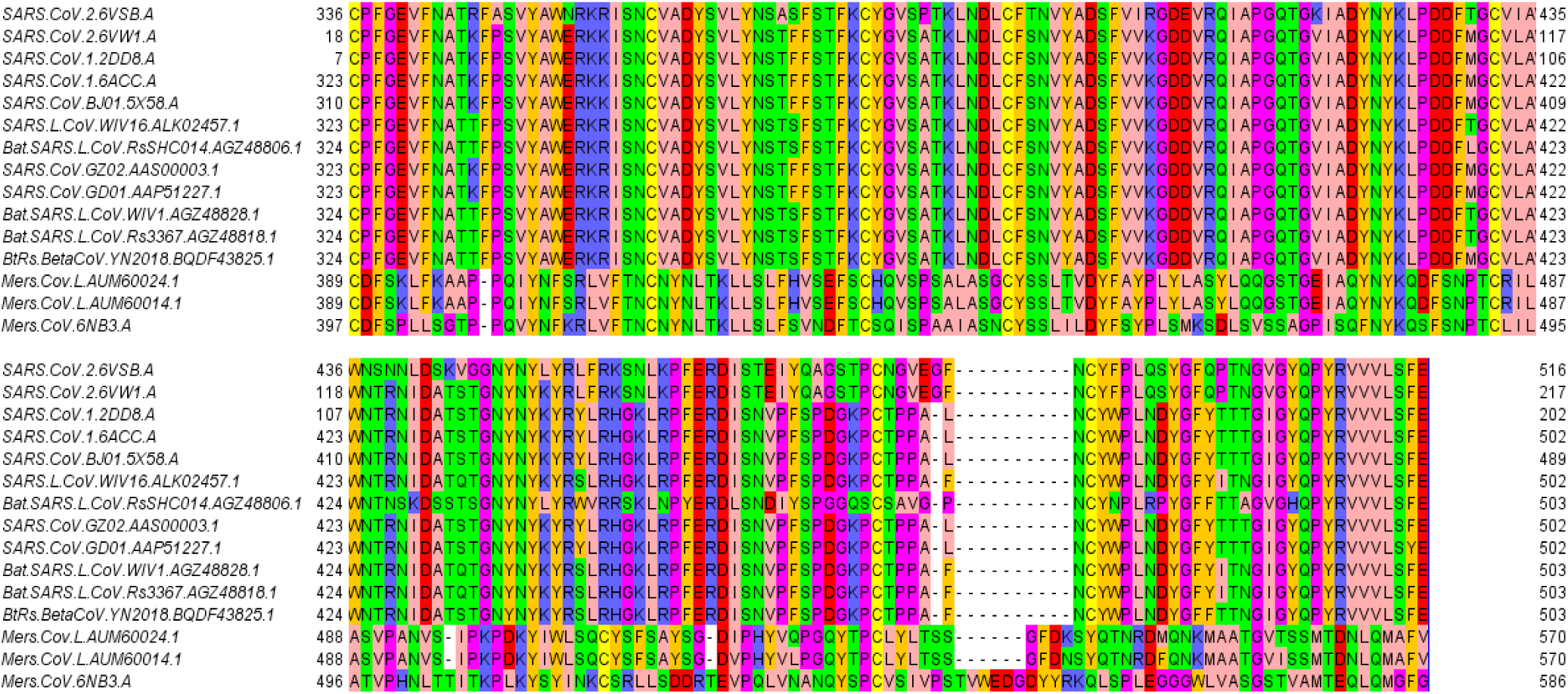
Multiple sequence alignment of RBDs from 11 SARS-CoV and 3 MERS-CoV strains. The reported residues numbering refers to the indicated sequences sampled by blastp or to the indicated crystallized structure sequences.

**Fig. 4.**
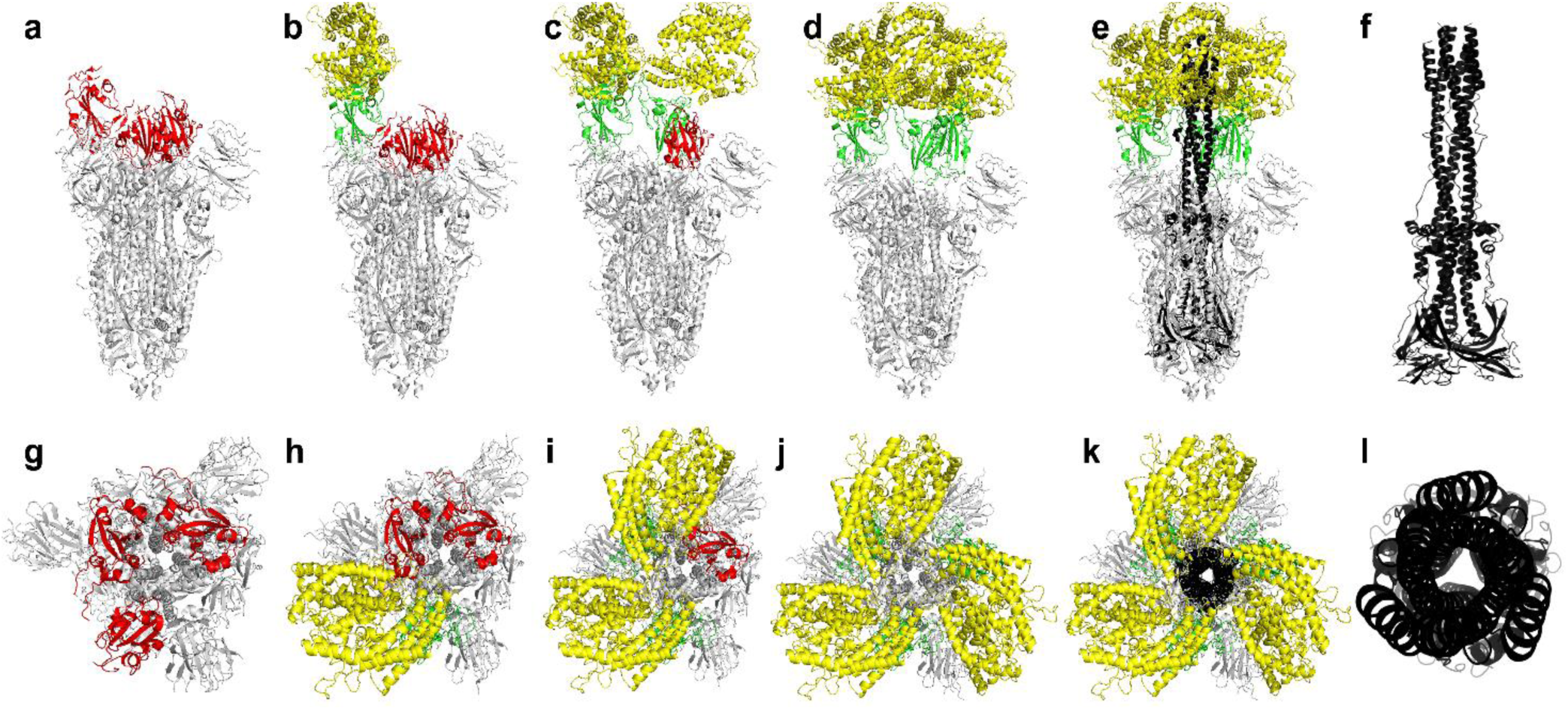
Side view (panels a-f) and top view (panes g-l) of H. sapiens SARS-CoV-2 spike protein interacting with 3 units of the human ACE2 N-terminal domain (panels a-e; g-k). SARS-CoV-2 spike protein trimer (6vsb.pdb) is reported in white cartoon representation with the 3 spike receptor binding domains reported in red (in the closed pre-fusion state) or green (in the open pre-fusion state) cartoon. The open pre-fusion state allows establishing pre-invasion interactions with ACE2 N-terminal domain. SARS-CoV-2 spike protein trimer C-terminal domain, resulting from protein cleavage that triggers the post-fusion conformation, is reported in black cartoon representation in panel f (lateral view) and l (top view).

It was possible to superimpose the crystallized SARS-CoV-2 spike protein in pre-fusion conformation and the modelled SARS-CoV-2 spike protein trimers in post-fusion conformation for showing the deep conformational changes occurring along conformation interconversion (Fig. 4). Apparently, the post-fusion conformation appears to be elongated and narrower than the pre-fusion conformation. The top portion of the post-fusion conformation locates beyond ACE2 receptors (Fig. 4), known for being anchored to plasma membrane and involved in internalization events (7, 41, 44, 45, 52).

### 3.3. SARS-CoV-1 and SARS-CoV-2 RBD residues involved in direct interactions with ACE2

From the available crystallized structures and from the obtained 3D structure models it was possible to highlight SARS-CoV-1 spike RBD (2ajf.pdb) and SARS-CoV-2 spike RBD residues (6) involved in the binding of the human ACE2 (Fig. 5 and Supp. Tab. 1). Notably, ion pair interactions observed between SARS-CoV-1 spike RBD and the human ACE2 are also observed between SARS-CoV-2 spike RBD and the human ACE2. The reported data represents an updated/integrated analysis of a similar ones reported in (53), in light of the recently deposited SARS-CoV-2 spike RBD in complex with the human ACE2 (6vw1.pdb).

**Fig. 5.**
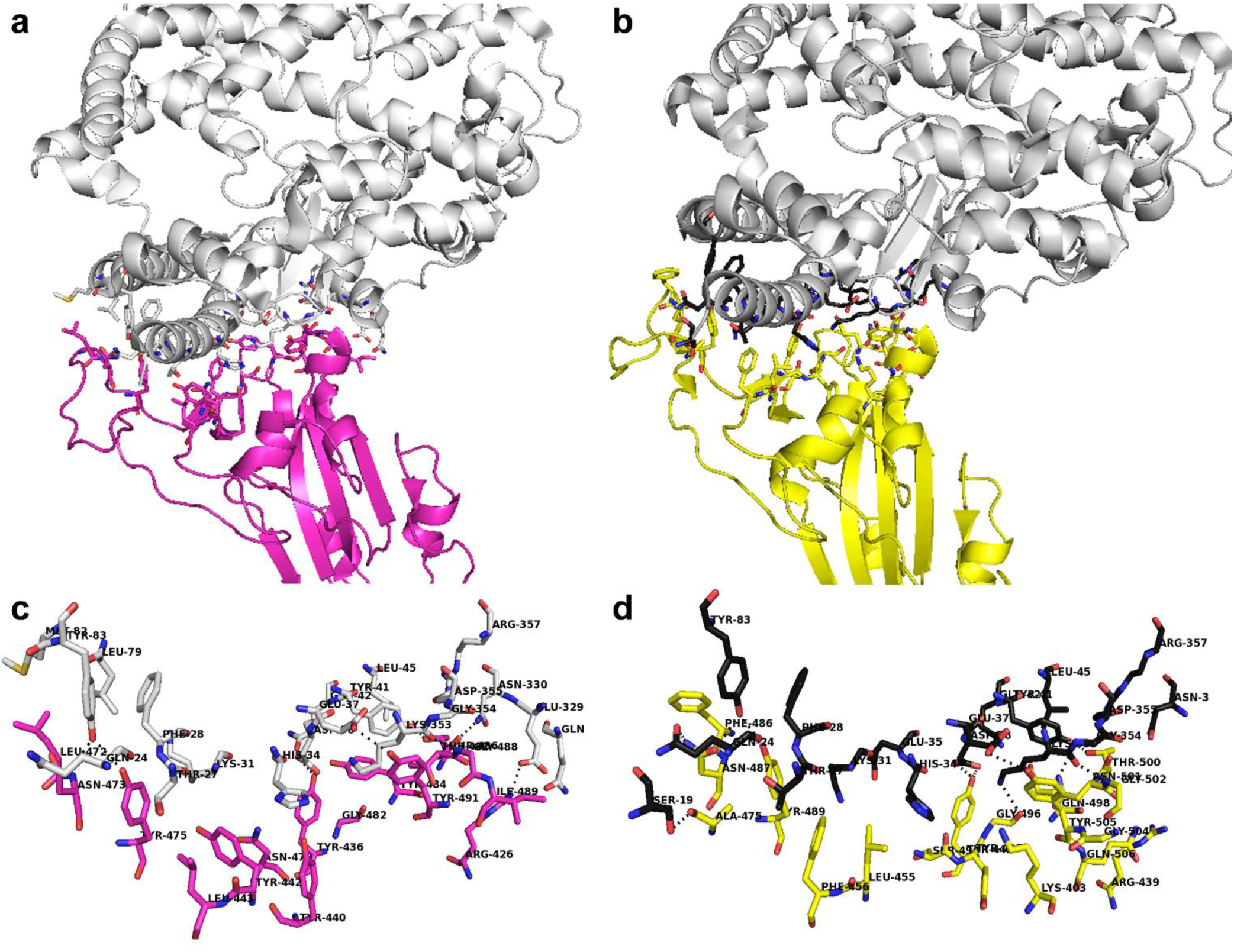
SARS-CoV-1 and SARS-CoV-2 RBD residues involved in direct interactions with ACE2. H. sapiens ACE2 is reported in white cartoon representation. SARS-CoV-1 RBD is reported in magenta cartoon representation, whereas SARS-CoV-2 RBD is reported in yellow cartoon representation. Panel a, c. Residues involved in polar interactions between SARS-CoV-1 RBD (magenta sticks) and ACE2 (white sticks). Panels b, d. Residues involved in polar interactions between SARS-CoV-2 RBD (yellow sticks) and ACE2 (black sticks). Polars interactions are represented by black dashed lines in the exploded views reported in panels c and d.

### 3.4. Comparative analysis of existing SARS.CoV.1. Spike RBD directed neutralizing antibodies and interaction predictions with SARS.CoV.2 Spike RBD

RBD from SARS-CoV-1 was crystallized in complex with the FAB domain of two different antibodies, namely m396 (2dd8.pdb, (44)) and S230 (6nb7.pdb, (45)). Both of them show high affinity for SARS-CoV-1 spike RBD, being able to block attachment to ACE2 (44, 45). Nevertheless, they show different peculiarities in their mechanism of action.

Indeed, S230 after binding RBD, similarly to ACE2, is able to trigger the SARS-CoV spike transition to the post-fusion conformation and it is not clarified yet, if virus-cell fusion may be triggered by S230 also when S230-RBD interactions occurs close to the surface of the cells target of the SARS-CoV-1 (45). At variance with S230, m396 antibody appears to be able to prevent SARS-CoV-1 spike-ACE2 interactions and SARS-CoV-1 spike pre-/post-fusion conformation transition, neutralizing virus attack (44).

Thanks to the high percentage of identical residues (> 75 %) between SARS-CoV-1 and SARS-CoV-2 spike RBD domains and to their highly similar tertiary structure, as observed from the RMSD of 0.5 Å between the coordinates of RBDs from SARS-CoV-1 (6nb7.pdb, (45) and 2dd8.pdb, (44)) and SARS-CoV-2 (6vw1.pdb (54) and 6vsb.pdb, (21)) spike proteins, it was possible to evaluate interactions between m396 and SARS-CoV-2 spike RBD and to propose a sequence/structure of an ideal FAB m396-based chimeric antibody for targeting SARS-CoV-2 spike RBD domain, preventing fusion events with ACE2 and thus the following infection.

With this aim, we firstly highlighted the different RBD portions bound to the known antibodies. Then, we superimposed SARS-CoV-1 RBD to SARS-CoV-2 RBD for highlighting differences in residues involved in direct interactions with m396 CDR regions and with S230 CDR regions (Fig. 6 and Tab. 1).

**Tab. 1.** List of SARS.CoV.1 and SARS.CoV2 RBD residues and of ACE2. Bold black residues delimited by borders indicate a pair or a cluster of residues involved in polar inter-protein interactions. Normal black residues indicate residues at the Spike-RBD.vs.ACE2 protein interface distant less than 4 Å. The longest chains were chosen within those crystallized structures with multiple chains, for highlighting the listed interacting residues.

| ACE2.interacting.residues.with<br>SARS.CoV.1.RBD (2ajf.pd)b |  | ACE2.interacting.residues.with<br>SARS.CoV.2.RBD (6vw1.pdb) |  |
| --- | --- | --- | --- |
| ACE2<br>(chain A) | SARS.CoV.1.RBD<br>(chain E) | ACE2<br>(Chain B) | SARS.CoV.2.RBD<br>(Chain F) |
|  |  | **S19** | **A475** |
| **Q24**<br>**Y83** | **N473**<br>**Y475** | **Q24**<br>**Y83** | **N487**<br>**Y489** |
|  |  | **E37** | **Y505** |
| **D38**<br>**Q42** | **Y436** | **D38** | **Y449** |
| **Y41**<br>**N330** | **T486** | **Y41** | **T500**<br>**N501** |
| **K353** | **T487** | **K353**<br>**G354** | **G496**<br>**G502** |
| **E329** | **R426** |  |  |
| T27 | L45 | T27 | K403 |
| F28 | Y83 | F28 | R439 |
| K31 | Y440 | K31 | L455 |
| H34 | Y442 | H34 | F456 |
| E37 | L443 | E35 | F486 |
| L45 | L472 | Q42 | S494 |
| L79 | N479 | L45 | Y495 |
| M82 | G482 | N330 | Q498 |
| Q325 | Y484 | D355 | G504 |
| N330 | G488 | R357 | Q506 |
| G354 | I489 |  |  |
| D355 | Y491 |  |  |
| R357 |  |  |  |

**Fig. 6.**
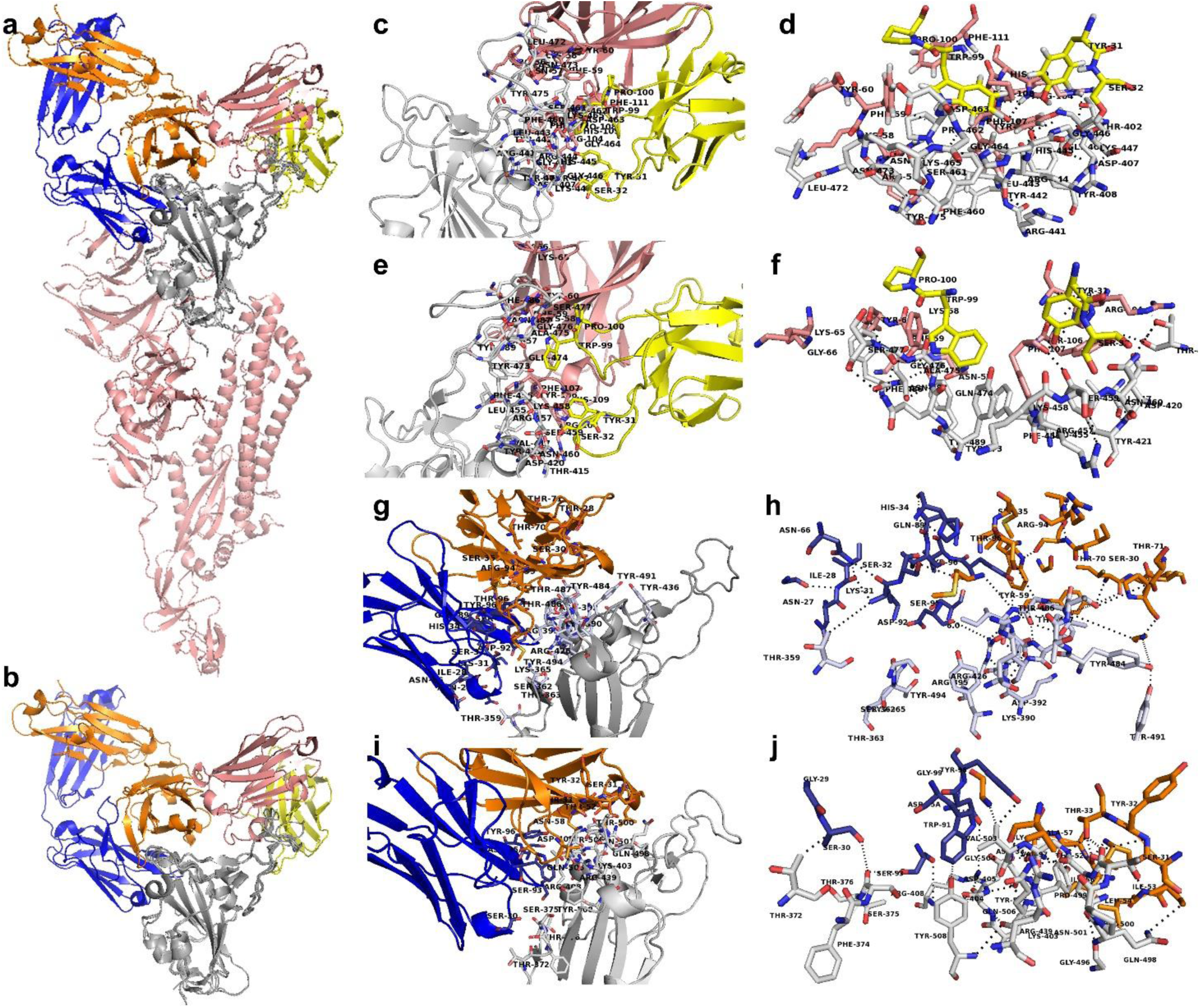
SARS.CoV.1 Spike and SARS.CoV.2 Spike monomers in pre-fusion conformation interacting with SARS.CoV.1 Spike RBD selective antibodies S230 (6nb7.pdb) and m396 (2dd8.pdb). Panel a. Superimposition of the tertiary structure of SARS.CoV.1 (6nb7.pdb) and SARS.CoV.2 (6vsb.pdb) spike protein monomers reported in pink cartoon representation. SARS.CoV.1 and SARS.CoV.2 RBDs are reported in grey cartoon representation. S230 FAB ab portion (6nb7.pdb) is reported in yellow (light chain) and pink (heavy chain) cartoon representation. m396 FAB ab portion (2dd8.pdb) is reported in orange (light chain) and blue (heavy chain) cartoon representation. Panel b. zoomed view of the superimposition of SARS.CoV.1 Spike and SARS.CoV.2 Spike RBD domains interacting with S230 and m396 FAB antibodies (see Panel a. for colors). Panel c-d. Super zoomed and rotated views of the crystallized SARS.CoV.1 Spike RBD residues interacting with S230 ab. Panel e-f. Super zoomed and rotated views of SARS.CoV.2 Spike RBD predicted residues interacting with S230 ab. Panel g-h. Super zoomed and rotated views of the crystallized SARS.CoV.1 Spike RBD residues interacting with m396 ab. Panel i-j. Super zoomed and rotated views of SARS.CoV.2 Spike RBD predicted residues interacting with m396 ab. Panels c-j Residues at the RBD – ab interface in the 3.5-4 Å distance range are reported in sticks representation. White sticks indicate RBD residues; orange and blue sticks indicate m396 ab residues, yellow and pink sticks indicate S230 ab residues.

### 3.5 SARS.CoV.2 Spike RBD directed neutralizing antibody engineering

Due to the uncertain data concerning fusion events and mechanism of action of S230 antibody, we built a new SARS-CoV-1/2 RBD directed antibody starting from the analysis of monomer-monomer interface interactions observed between the m396 antibody crystallized in complex with SARS-CoV-1 RBD (44), superimposed to SARS-CoV-1 spike RBD / ACE2 complex (2ajf.pdb), and by comparing them with monomer-monomer interface interactions observed between the modelled m396 antibody in complex with SARS-CoV-2 RBD, superimposed to SARS-CoV-2 RBD/ACE2 (6vw1.pdb; 6lzg.pdb) protein complex (Fig. 7).

**Fig. 7.**
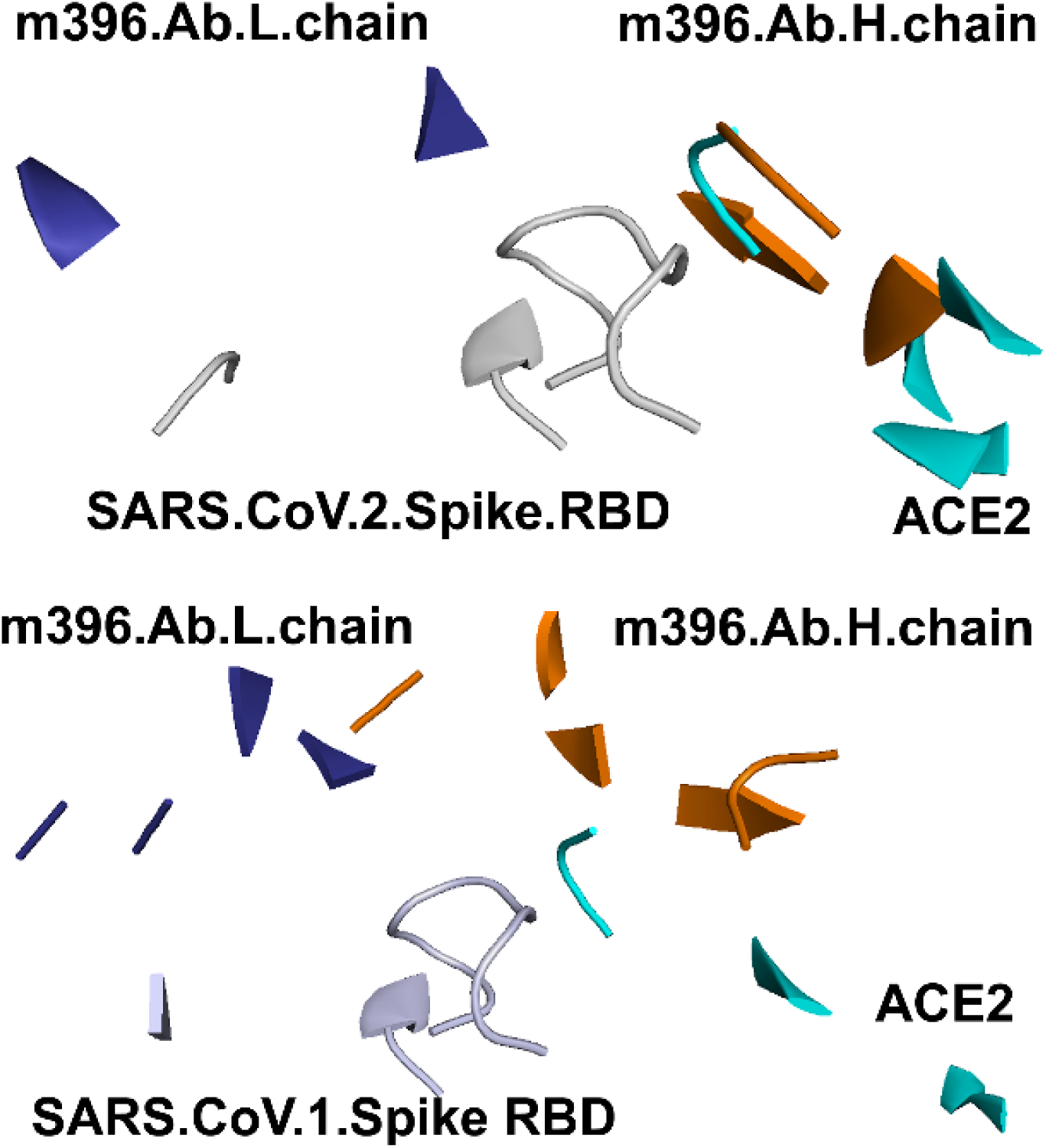
Molecular framework of the investigated proteins hosting SARS-CoV-spike RBDs, light and heavy chain of the m396 antibody and the human ACE2, simultaneously. The shown spike RBD, ACE2 and m396 protein portions are those in a reciprocal distance range of 4 Å. Upper panel: Superimposition of the crystallized SARS-CoV-1 spike RBD (white cartoon representation) in complex with m396 antibody (2d88.pdb, orange and blue cartoon) and ACE2 (2ajf.pdb, cyan cartoon). Bottom panel: superimposition of SARS-CoV-2 spike RBDs (from 6vw1.pdb, white cartoon representation), ACE2 from 6vw1.pdb (cyan cartoon) and m396 from 2d88.pdb (orange and blue cartoon).

Then, we highlighted m396 CDR residues (Tab. 2) for replacing them aiming to increase m392 affinity versus SARS-CoV-1/2 spike RBDs. Residues to be mutated/replaced were chosen according to space-restraints and chemical needs for better complementing SARS-CoV-1/2 spike RBD surface, based on the available SARS-CoV-1/2 RBD structures in complex with ACE2, aiming to produce something that resembled ACE2 surface (Tab. 1-2). Some of the proposed mutated residues (Tab. 3) are surely allowed because already observed at the corresponding sites of other known antibodies, according to Chotia/Kabat rules (http://www.bioinf.org.uk/abs/chothia.html; (55)). Residues replacement was directly performed in the newly generated 3D model hosting the interacting m396-SARS-CoV-2 spike RBD. Similarly, a complex of the modified m396 antibody interacting with SARS-CoV-1 RBD was also created. All m396 CDR mutated residues are reported in Tab. 3. Furthermore, mutated residues within m396 CDR interacting with SARS-CoV-2 spike RBD residues can be observed in Fig. 8.

**Tab. 2.**
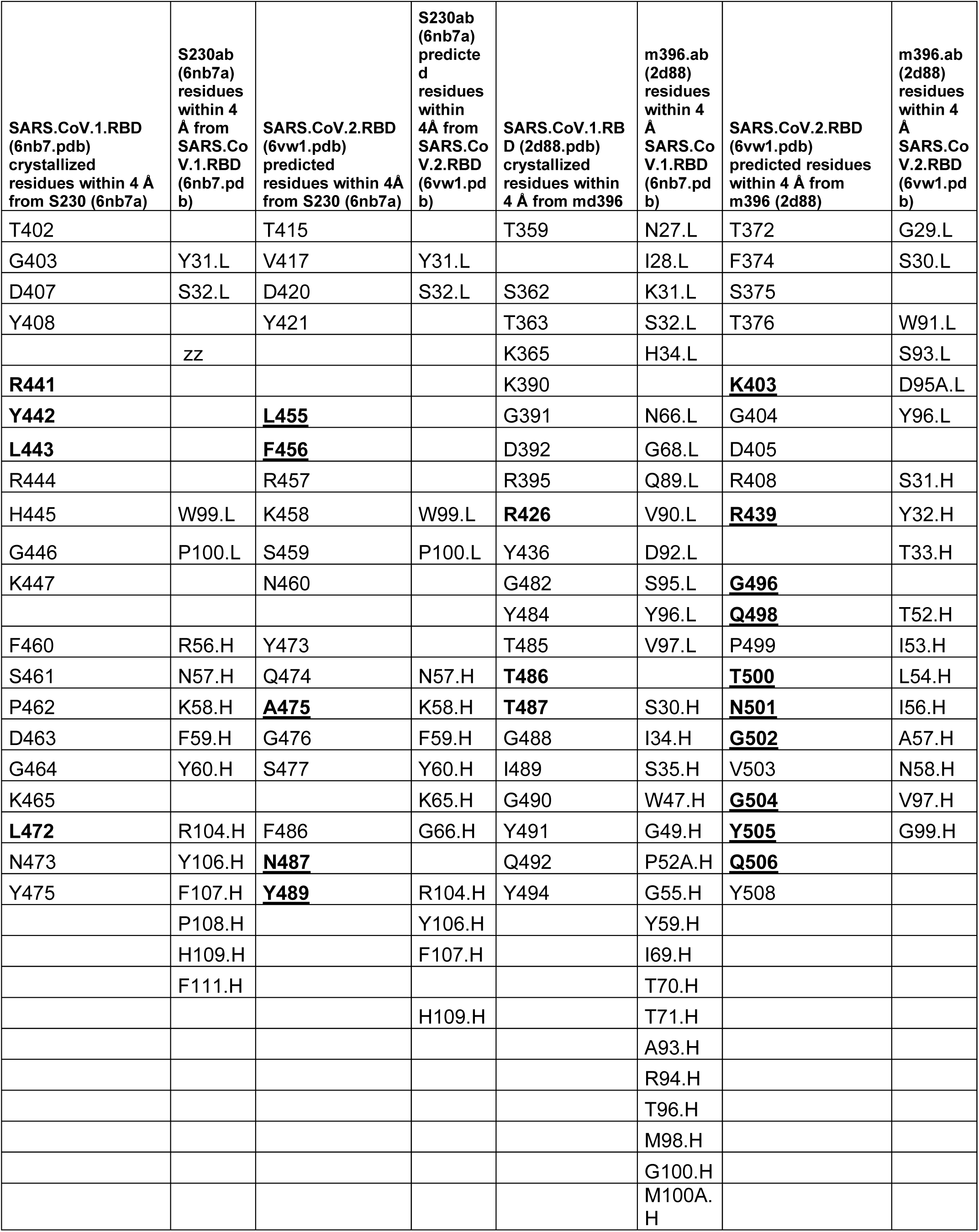
List of SARS-CoV-1/2 RBD residues within 4 Å from S230/m396 antibody residues. Bold residues indicate SARS.CoV.1 residues interacting alternatively with both ACE2 and/or m396/S230 in the crystallized available structures. Distance range below 4 Å. Bold underlined residues indicate SARS.CoV.2 residues interacting with ACE2 and predicted to interact with m396 in a distance range below 4 Å

| SARS.CoV.1.RBD<br>(6nb7.pdb)<br>crystallized<br>residues within 4 Å<br>from S230 (6nb7a) | S230ab<br>(6nb7a)<br>residues<br>within 4<br>Å from<br>SARS.Co<br>V.1.RBD<br>(6nb7.pd<br>b) | SARS.CoV.2.RBD<br>(6vw1.pdb)<br>predicted<br>residues within 4Å<br>from S230 (6nb7a) | S230ab<br>(6nb7a)<br>predicte<br>d<br>residues<br>within<br>4Å from<br>SARS.Co<br>V.2.RBD<br>(6vw1.pd<br>b) | SARS.CoV.1.RB<br>D (2d88.pdb)<br>crystallized<br>residues within<br>4 Å from md396 | m396.ab<br>(2d88)<br>residues<br>within 4<br>Å<br>SARS.Co<br>V.1.RBD<br>(6nb7.pd<br>b) | SARS.CoV.2.RBD<br>(6vw1.pdb)<br>predicted residues<br>within 4 Å from<br>m396 (2d88) | m396.ab<br>(2d88)<br>residues<br>within 4<br>Å<br>SARS.Co<br>V.2.RBD<br>(6vw1.pd<br>b) |
| --- | --- | --- | --- | --- | --- | --- | --- |
| T402 |  | T415 |  | T359 | N27.L | T372 | G29.L |
| G403 | Y31.L | V417 | Y31.L |  | I28.L | F374 | S30.L |
| D407 | S32.L | D420 | S32.L | S362 | K31.L | S375 |  |
| Y408 |  | Y421 |  | T363 | S32.L | T376 | W91.L |
|  | zz |  |  | K365 | H34.L |  | S93.L |
| **R441** |  |  |  | K390 |  | **K403** | D95A.L |
| **Y442** |  | **L455** |  | G391 | N66.L | G404 | Y96.L |
| **L443** |  | **F456** |  | D392 | G68.L | D405 |  |
| R444 |  | R457 |  | R395 | Q89.L | R408 | S31.H |
| H445 | W99.L | K458 | W99.L | **R426** | V90.L | **R439** | Y32.H |
| G446 | P100.L | S459 | P100.L | Y436 | D92.L |  | T33.H |
| K447 |  | N460 |  | G482 | S95.L | **G496** |  |
|  |  |  |  | Y484 | Y96.L | **Q498** | T52.H |
| F460 | R56.H | Y473 |  | T485 | V97.L | P499 | I53.H |
| S461 | N57.H | Q474 | N57.H | **T486** |  | **T500** | L54.H |
| P462 | K58.H | **A475** | K58.H | **T487** | S30.H | **N501** | I56.H |
| D463 | F59.H | G476 | F59.H | G488 | I34.H | **G502** | A57.H |
| G464 | Y60.H | S477 | Y60.H | I489 | S35.H | V503 | N58.H |
| K465 |  |  | K65.H | G490 | W47.H | **G504** | V97.H |
| **L472** | R104.H | F486 | G66.H | Y491 | G49.H | **Y505** | G99.H |
| N473 | Y106.H | **N487** |  | Q492 | P52A.H | **Q506** |  |
| Y475 | F107.H | **Y489** | R104.H | Y494 | G55.H | Y508 |  |
|  | P108.H |  | Y106.H |  | Y59.H |  |  |
|  | H109.H |  | F107.H |  | I69.H |  |  |
|  | F111.H |  |  |  | T70.H |  |  |
|  |  |  | H109.H |  | T71.H |  |  |
|  |  |  |  |  | A93.H |  |  |
|  |  |  |  |  | R94.H |  |  |
|  |  |  |  |  | T96.H |  |  |
|  |  |  |  |  | M98.H |  |  |
|  |  |  |  |  | G100.H |  |  |
|  |  |  |  |  | M100A.<br>H |  |  |

**Tab. 3.** List of CDR L/H residues detectable in m396 antibody according to Chotia/Kabat classification. The investigated (*) built variants (column “Mutations based on space restraints needs”) and (#) known mutations (column “Allowed variants”) according to Chotia/Kabat rules are also reported for comparative purposes.

|  | Mutations based on space restraints needs* | Allowed variants# |  | Mutations based on space restraints needs* | Allowed variants# |
| --- | --- | --- | --- | --- | --- |
| **CDR-L1** |  |  | **CDR-H1** |  |  |
| 24-GGNNIGSKSVH-34 | S30R;<br>K31R | G25A;<br>N26S;<br>I28N/S/D/E;<br>G29I/V;<br>S32Y | 26-GGTFSSYTIS-35 | S31K;<br>T33E |  |
| **CDR-L2** |  |  | **CDR-H2** |  |  |
| 50-DDSDRPS-56 |  | D51A/T/G/V | 50-GITPILGIANYAQKFQG-66 | L55D;<br>I57Y; | T52D/L/N/S/Y;<br>I54A/G/Y/S/K/T/N;<br>L55N/S/T/K/D/G;<br>I57Y/R/E/D/G/V/S/A |
| **CDR-L3** |  |  | **CDR-H3** |  |  |
| 89-QVWDSSSDYV-98 | S94E;<br>S95R | V90Q;<br>D92S; Y97I;<br>V98T | 99-DTVMGGMDV-17 | V101K;<br>G103L |  |

**Fig. 8.**
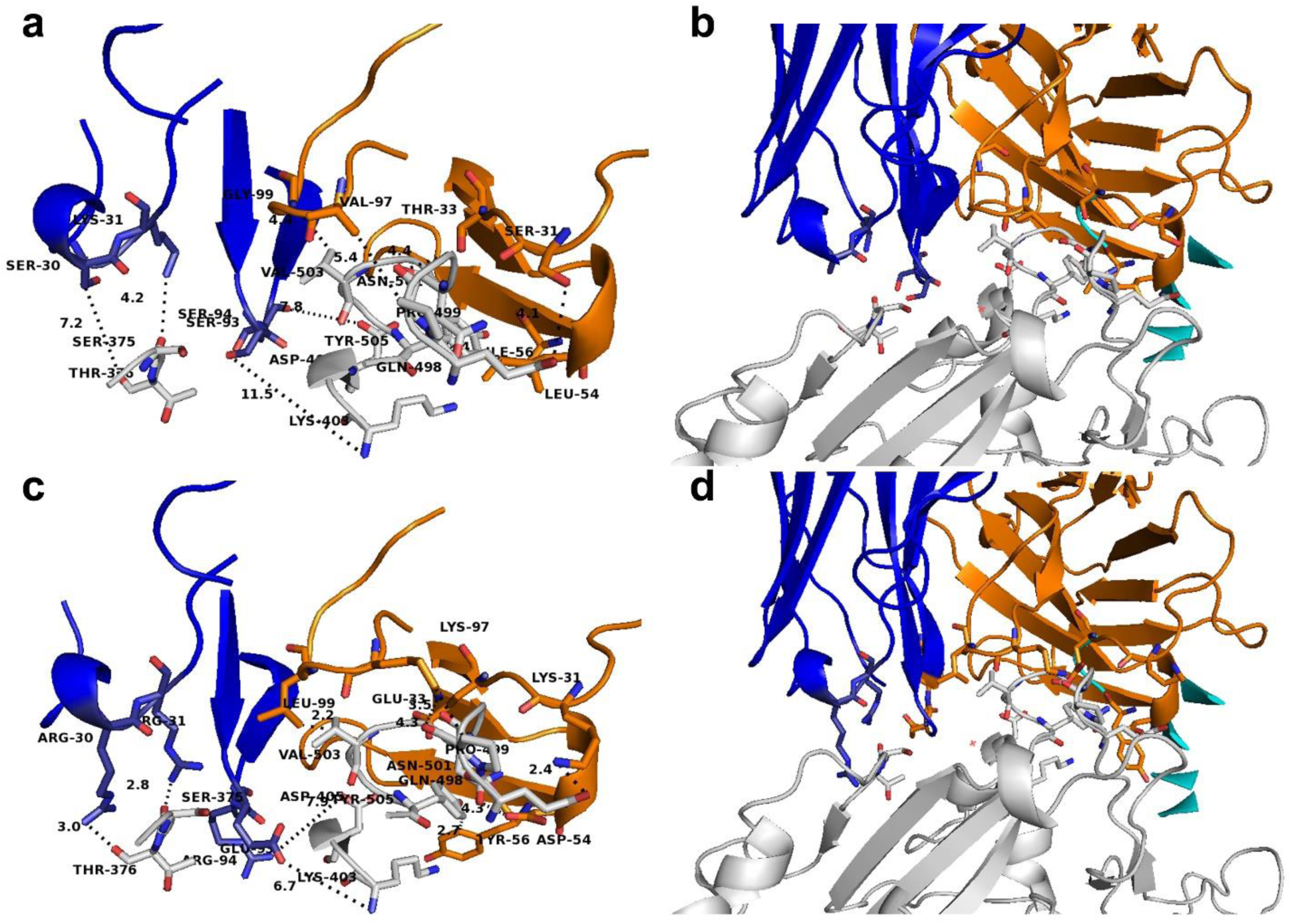
m396 neutralizing antibody, native and engineered, in complex with SARS-CoV-2 spike RBD. Panels a-b: exploded view and perspective view of native m396 neutralizing antibody (in orange blue cartoon) in complex with SARS-CoV-2 spike RBD (in white cartoon representation). Residues at the m396/RBD interface in a distance range withni 4 are indicated by white sticks (RBD), orange sticks (m396 CDR-H residues) and blue sticks (m396 CDR-L residues). Panels c-d: exploded view and perspective view of the engineered m396 predicted neutralizing antibody (in orange blue cartoon) in complex with SARS-CoV-2 spike RBD (in white cartoon representation). Residues at the engineered m396/RBD interface in a distance range within 4 Å are indicated by white sticks (RBD), orange sticks (engineered m396 CDR-H residues) and blue sticks (engieered m396 CDR-L residues).

The engineered FAB portions were thus aligned and superimposed on the FAB portion of a crystallized IgG (1igt.pdb, (48)). The sequence of the chimeric antibodies can be observed in Supp. Fig. 1, whereas their complete structure can be observed in Supp. Fig. 2.

### 3.6 Free energy calculation

The interaction energies calculated between the SARS-CoV-2 spike RBD domain and m396 native antibody FAB portion gives a negative value (Tab. 4), confirming that there might be a binding interaction between m396 native antibody FAB portion and SARS-CoV-2 spike RBD. This result is encouraging, also due to the indirect validation obtained by getting similar interaction energies for the crystallized SARS-CoV-1 RBD in complex with m396 (2d88.pdb) and for SARS-CoV-1 and SARS-CoV-2 spike RBD domains crystallized in complex with ACE2 (2ajf.pdb and 6vw1.pdb, respectively) (Tab. 4). Furthermore, a strong interaction (in terms of interaction energies calculated by FoldX Analyse complex assay) is also predicted between SARS-CoV-2 spike RBD (but also SARS-CoV-1 spike RBD) and the modified m396 antibody (see Tab. 4), suggesting that the engineered m396 might be more efficient than the native m396 in binding the SARS-CoV-2 (more than SARS-CoV-1) spike RBD.

**Tab. 4.**
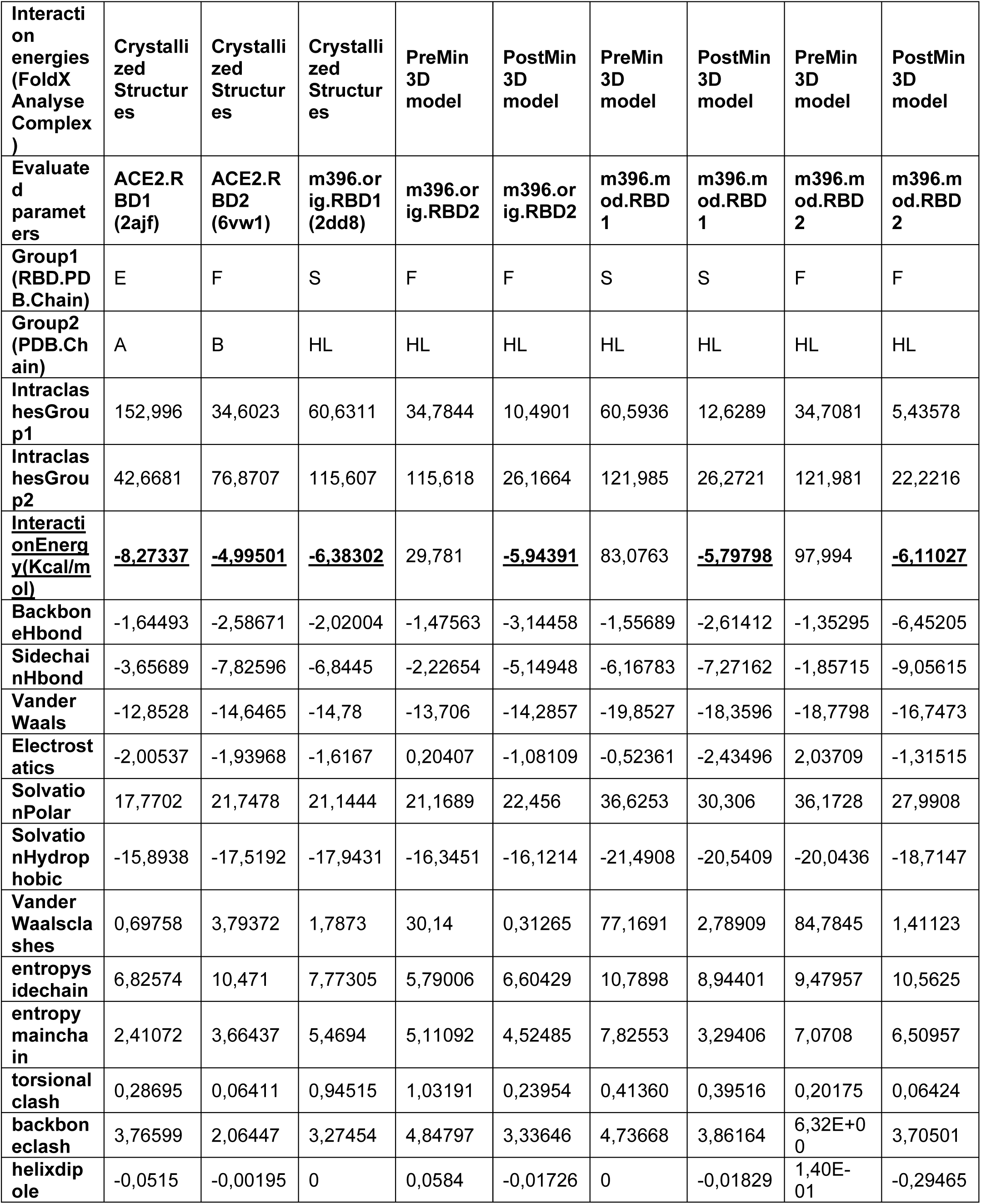
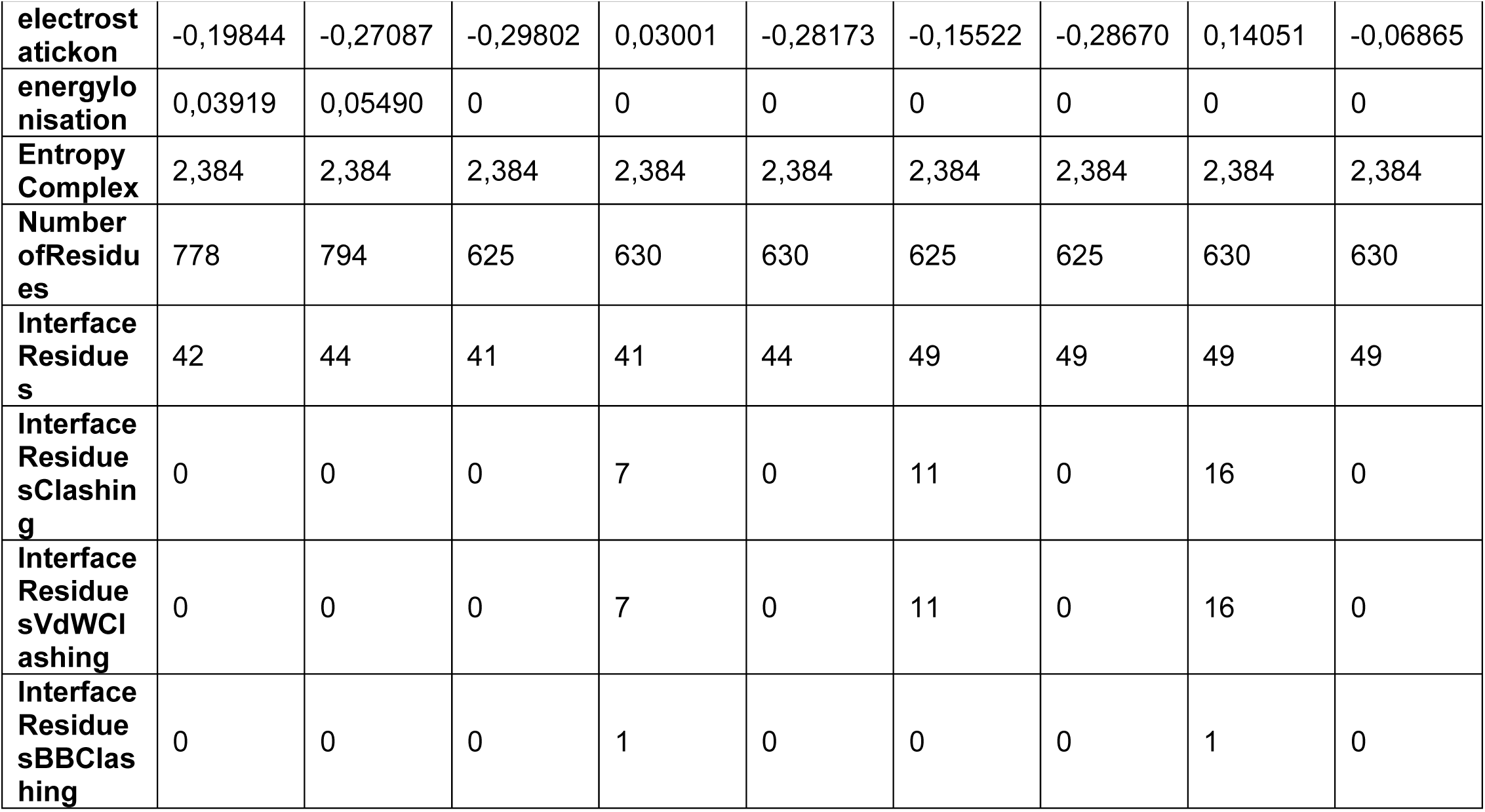
Energy calculations on crystallized structures or 3D comparative models of the investigated protein complex. The PDB.Chain indicates the chain of the PDB used within the indicated analyses on the cited crystallized structures or models obtained by superimposition with the indicated chains. The longest chains were chosen for the “interaction energy” analyses for those crystallized structures with multiple chains. Chain E, F, and S indicate the RBD chain within the investigated PDB_IDs. Chain A and B indicate the ACE2 chain within the investigated PDB_IDs. Chain H, L indicate the heavy and light chain of the investigated antibody (wild type and engineered variants), according to the indicated PDB_ID. PreMin and PostMin refer to models prior and after energy minimization performed on the Yasara minimization server.

## DISCUSSION

The indicated pipeline has allowed to set-up a molecular framework hosting SARS-CoV-2 spike protein, ACE2 receptor and different antibodies in the same pdb session that could be handled with different molecular visualizers. In this molecular framework it is possible to study and predict, at molecular level, interactions between the different “pieces” of the framework that my help in understanding virus invasion mechanisms, developing new vaccines or antibodies, identifying small molecules with high affinity for viral proteins and establishing quick/safe diagnosis selective/specific kits. Indeed, the scientific community is now focused in the development of new weapons for containing SARS-CoV-2 spread and COVID-19 complications as it could be observed in the enormous effort in developing new vaccines based on a virus protein/nucleic acid portion able to induce an efficient and specific immunogenic response(56–60), or in developing a neutralizing antibody highly specific for SARS-CoV-2 spike RBD (25, 26, 61–64), or in identifying chemicals with high affinity for SARS-CoV-2 crucial proteins (1–3, 65–67).

Within the presented molecular framework, we have highlighted a set of possible efficient interactions between the crystallized m396 antibody and SARS-CoV-2 spike RBD, raising the question about the possibility to test directly m396 on cultured cells exposed to the virus and then, hopefully, on patients.

Starting from that observation we have also proposed a set of modifications of m396 CDR residues resulting in a higher specific antibody, to be expressed and tested on cultured cells. Along the development of our antibody engineering modeling session an important paper was published and another is under revision in support of the hypothesis that m396 may be able to bind SARS-CoV-2 spike protein (61, 62).

It was also possible to pose in the proposed molecular framework the recent proposed SARS-CoV-2 spike RBD directed CR3022 FAB antibody (6yla.pdb; 6w41.pdb, (62)), showing that it binds a different site of RBD that protrudes towards the central cavity of the spike protein trimer (data not shown). It appears that the RBD-antibody interaction is possible only if at least two RBDs on the trimeric spike protein are in the “open” state of the prefusion conformation and slightly rotated, in a site distant from ACE2 receptor binding region, according to what proposed by the authors (62). Dedicated studies are necessary for understanding if steric hindering problems might rise by using the whole antibody, and deepening the comprehension of the not competitive mechanism that would be observed between CR3022 and RBD in presence of ACE2 receptor.

Studying all the cited interactions in the same pdb-molecular session has allowed to highlight maybe the most crucial ACE2 portions involved in direct interactions with SARS-CoV-2 RBD, suggesting that the administration of the recombinant RBD, a spike monomer or the entire spike trimer, if correctly folded, might result in the efficient triggering of antibody production from our plasma b-cells, reducing COVID-19 complications (supporting what has been recently proposed (56–60)).

At the same time, the ACE2-RBD interactions estimated in our molecular framework has strengthened the hypothesis to use the recombinant ACE2 for limiting COVID-19 infection complications (according to what recently proposed (68, 69)).

A molecular framework like the ones here proposed will also help in studying the putative role of ACE inhibitors in perturbing ACE2-RBD interactions. Indeed, it was recently proposed that patients treated with ACE inhibitors might be more exposed to SARS-CoV-2 infection (70). Although ACE1 (refseq accession number: NP_068576.1, representing the main target of ACE inhibitors) and ACE2 (NP_690043.1, testis isoform or NP_000780, somatic isoform, among the most studied isoforms) share the 40 % of identical residues, few uncertain data about ACE inhibitors and a possible greater selectivity for ACE1 versus ACE2, or on their effect on ACE1/2 expression regulation are available in literature (70, 71). From a structural comparison it is observed that the RMSD of the crystallized native ACE2 coordinates (1r42.pdb, (72)) and ACE1 coordinates (1o8a.pdb, (73)) is lower than 2.5 Å.

Notably, the presence of ACE inhibitors captopril and enalaprilat (1uze.pdb,(74); 4c2p.pdb, (75)) and lisinopril (1o86.pdb, (73)) produces an RMSD lower than 0.3 Å in the atomic coordinates of the cited crystallized structures with reference to the native ACE1 (1r42.pdb, (72)).

Conversely, we cannot establish if the slightly higher RMSD observed between the native ACE2 (1r42.pdb) and ACE2 complexed with SARS-CoV-1 spike RBD (0. 41 Å, 2ajf.pdb) and SARS-CoV-2 spike RBD (1.2 Å, 6vw1.pdb) can be attributed exclusively to interactions with SARS-CoV-RBD, because the observed RMSDs are of the same order of magnitude of the experimental resolution of the investigated crystallized structures.

However, also admitting that ACE1 inhibitors at the employed dosage would target ACE2, with the same efficiency observed versus ACE1, the presence of those inhibitors in ACE2 binding cavity should not be able to induce an important conformational change in ACE2, which might favour a greater affinity of ACE2 versus SARS-CoV-2 spike RBD.

Thus, the only mechanism for which, patients treated with ACE inhibitors would be more exposed to SARS-CoV-2, would rely in a positive feedback induced by ACE inhibitors in ACE2 expression. Nevertheless, evidences in support of this hypothesis need to be deepened (71, 76).

In conclusion, the presented analysis highlights the importance to use fold recognition tools along the approach to a drug design problem according to a rational protocol (similar to what previously reported (29, 30, 49)), like the ones presented. Indeed, in this case, fold recognition tools have helped us in identifying crystallized structures of ACE2, SARS-CoV-spike proteins similar to those under investigation. Furthermore, performing structural comparative analysis has allowed to identify a possible good starting point, like the ones represented by m396, already crystallized in complex with SARS-CoV-1 spike RBD, for building the proposed antibodies. The same strategy might be applied also for future infections by those researchers involved in drawing new antibodies and/or developing new vaccines, i.e. for dealing with future coronaviruses.

To the best of our knowledge the reported SARS-CoV-2 spike protein trimer 3D model is the first model describing a possible conformational change leading to a reliable SARS-CoV-2 spike protein in post fusion conformation. The proposed model, based on the murine CoV spike protein (6b3o.pdb) crystallized in post fusion conformation, will help in understanding the mechanism allowing the virus envelop fusion with host cell plasma membranes, through and following interactions with ACE2.

Furthermore, the provided 3D model in post-fusion conformation according to the crystallized 3D structure of SARS-CoV-2 spike protein in pre-fusion conformation confirms the presence and stability of a sort of channel at the interface of the three monomers that could represent a good target site of a virtual screening of a chemical/drug library aiming to identify a small molecule/peptide with high affinity for a monomer (similarly to what proposed for EK1 peptide (77)), for preventing trimer formation and stabilization, or a small molecule/peptide with high affinity for the trimer, aiming to prevent conformational changes leading to the fusion of the viral envelope with host cell plasma membranes. The screening of a drug library would help in identifying an already approved drug with high affinity for the spike channel, that might be immediately tested on the bed-side, in the context of the drug-repositioning approaches (78, 79).

Notably, the provided molecular framework for investigating/drawing new antibodies based on space-restraints needs, would be used for the set-up of new antibodies based on the available tissue-specific immunoglobulin structures, as the proposed IgG2A (1igt.pdb, (48)) or other specialized antibodies, already optimized for targeting specific cells or receptors (i.e.1hzh.pdb, (80)), also among those that may successfully target the respiratory tract (1r70.pdb, (81) or 2qtj.pdb (82) or 6ue7.pdb (83)), that might be administered even by aerosol (84, 85).

At the same time, already at the preclinical level, the administered vaccines based on the administration of the entire SARS-CoV-2 spike protein ((56–58) or on the administration of the single SARS-CoV-2 spike RBD, will induce the production of specific antibodies that might be sequenced and modelled *in silico*. On this concern, the provided molecular network will help in quantifying interactions between SARS-CoV-2 RBD (also in cases of different RBD variants (86)) and the newly investigated antibodies, i.e.lower the calculated binding energy in the modelled complex, higher the likelihood to have more success with the investigated vaccines/antibodies.

The discovered antibodies with the highest affinity for RBD might also be implemented in a diagnosis kit aiming to the early identification of SARS-CoV-2 in sera, also in asymptomatic people.

Conversely, a new diagnosis kit could also be based on the native RBD or a modified synthetic RBD, with greater affinity for the detected human antibodies directed against SARS-CoV-2 spike RBD protein, for determining the real number of healthy people already exposed to the virus in the population.

The lacking knowledge about the real number of people exposed to the virus (including asymptomatic people, people with mild symptoms and rescued people that never needed hospitalization or quarantine) is the only important data that we still miss. Without data about the real number of people, exposed to the virus, in the population, coming back to normal life will be extremely slower.

### About technical questions

Detailed instructions for the set-up of the shown molecular framework have been provided in the manuscript. Nevertheless, we can also provide free assistance for academic analyses, upon request. We can also provide dedicated technical support for analyses requested by private companies through our BROWSer s.r.l. spin-off (in this case, please, write to and to CLP in Cc).

## ACKNOWLEDGEMENTS

Authors would like to thank the Italian Association for Mitochondrial Research (www.mitoairm.it), IT resources made available by ReCaS, a project funded by the MIUR (Italian Ministry for Education, University and Re-search) in the “PON Ricerca e Competitività 2007–2013-Azione I-Interventi di ra_orzamentostrutturale” PONa3_00052, Avviso 254/Ric, University of Bari (“Fondi Ateneo ex-60%”2016”; “ProgettoCompetitivo 2018” and “FFABR 2017-2018”). Authors would also like to thank MIUR for having funded the project “Salute, alimentazione, qualità della vita”: individuazione di un set di biomarker dell’apoptosi” for an innovative industrial PhD course—PON RI 2014-2020, CUP H92H18000160006. Authors would also like to thank Angelo Onofrio (Biotechnologist) and Luna Laera (Biotechnologist) for critical reading and/or stimulating discussions.

## Supporting Material

**Supp. Tab. 1.**
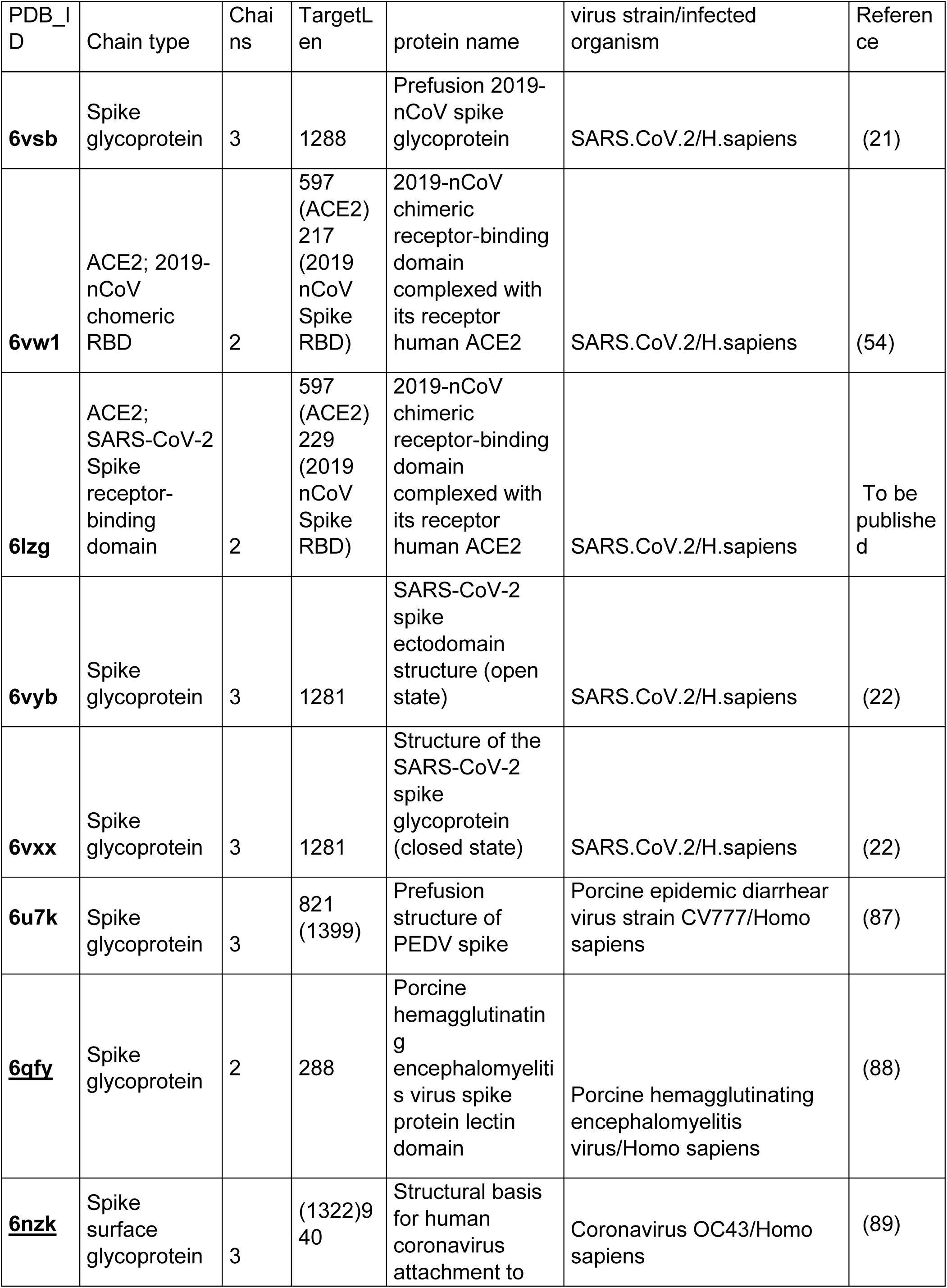
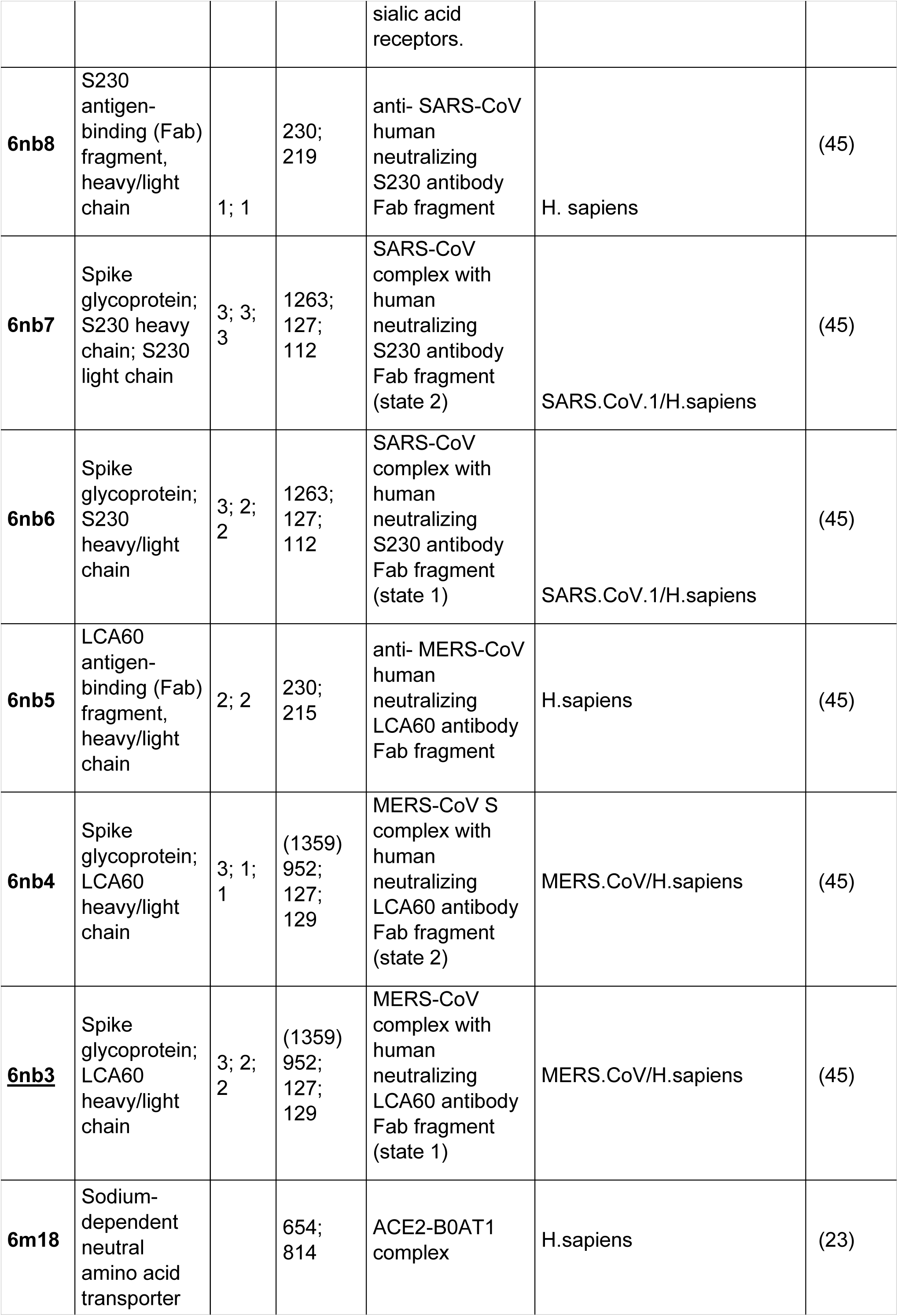
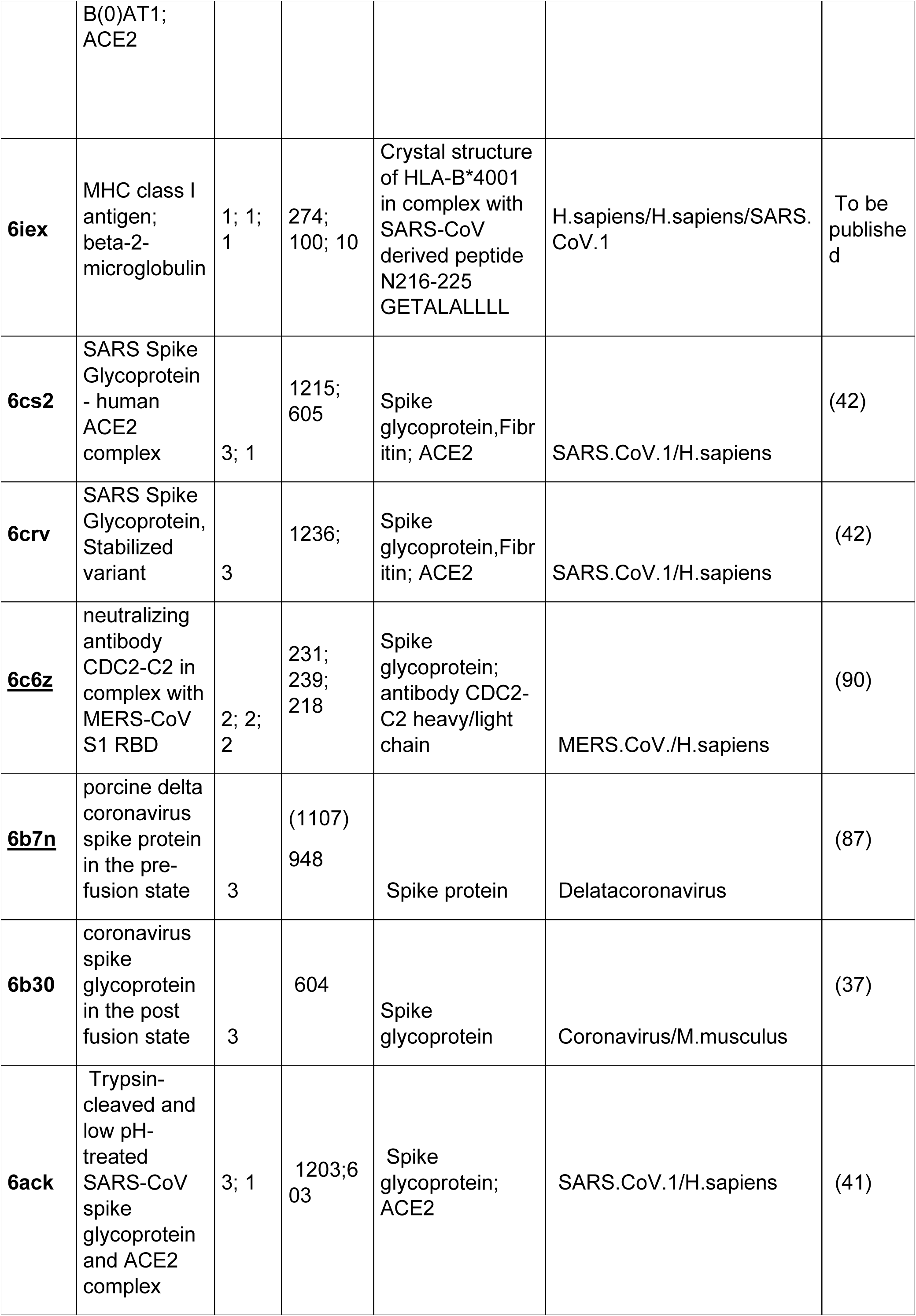
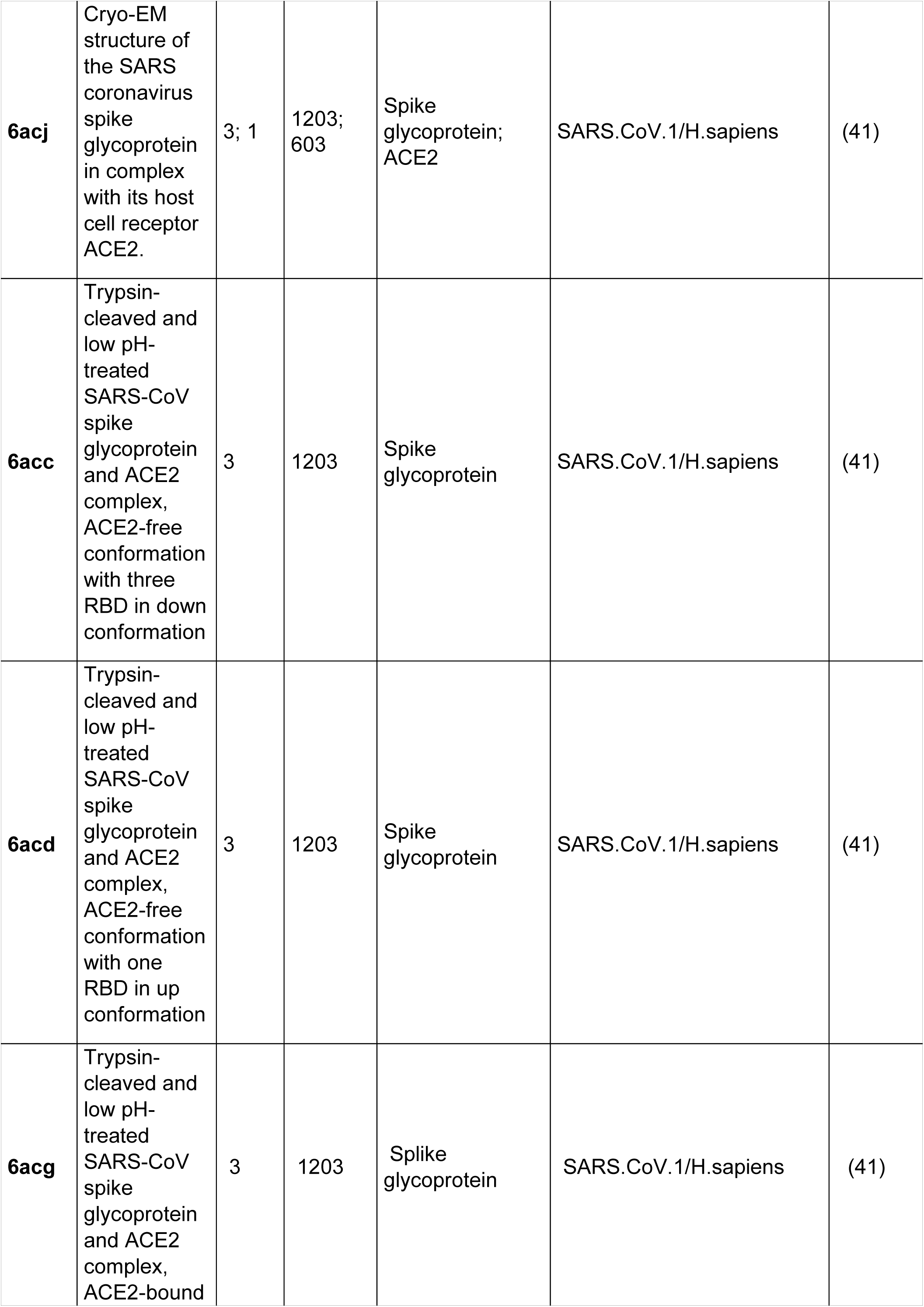
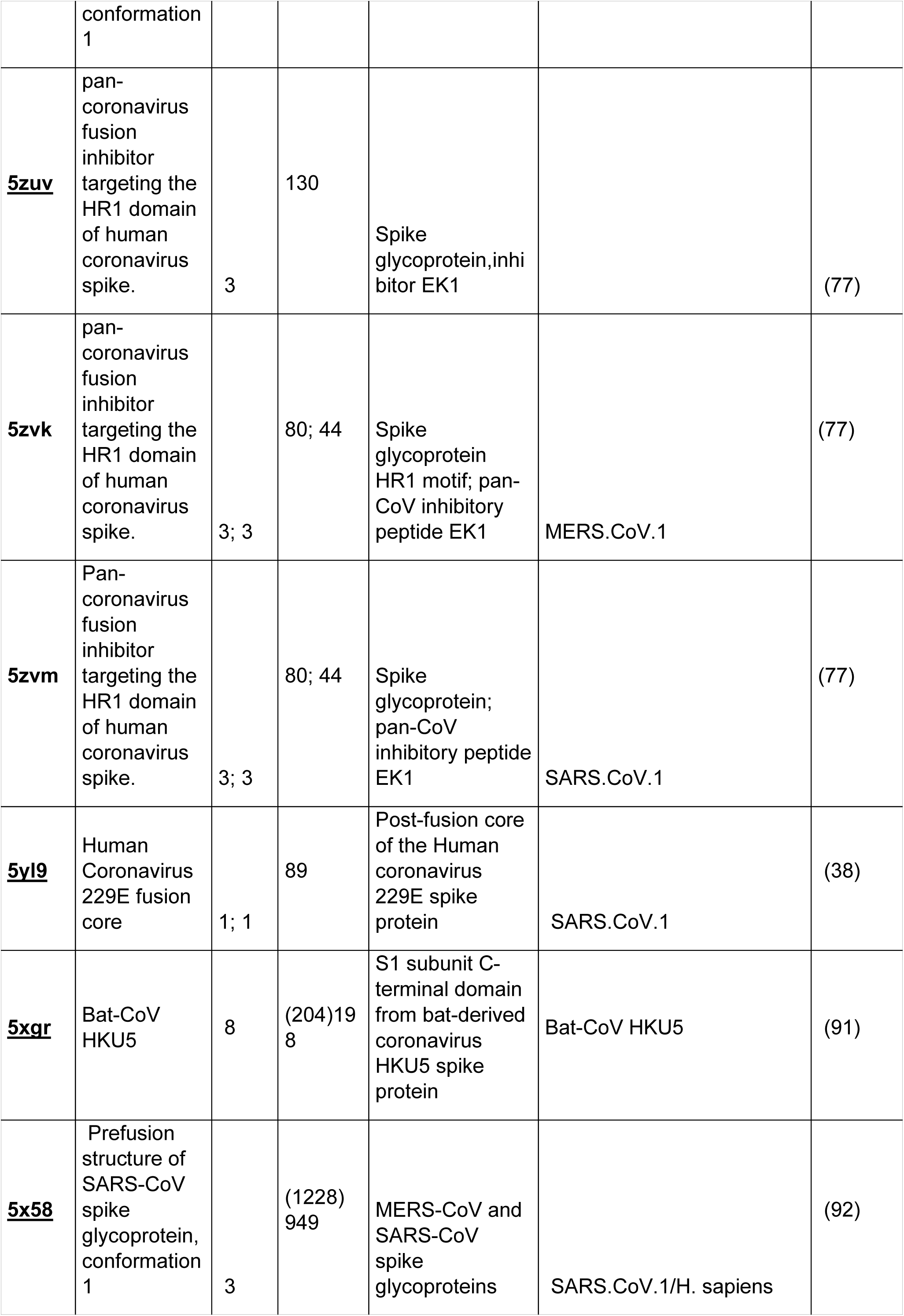
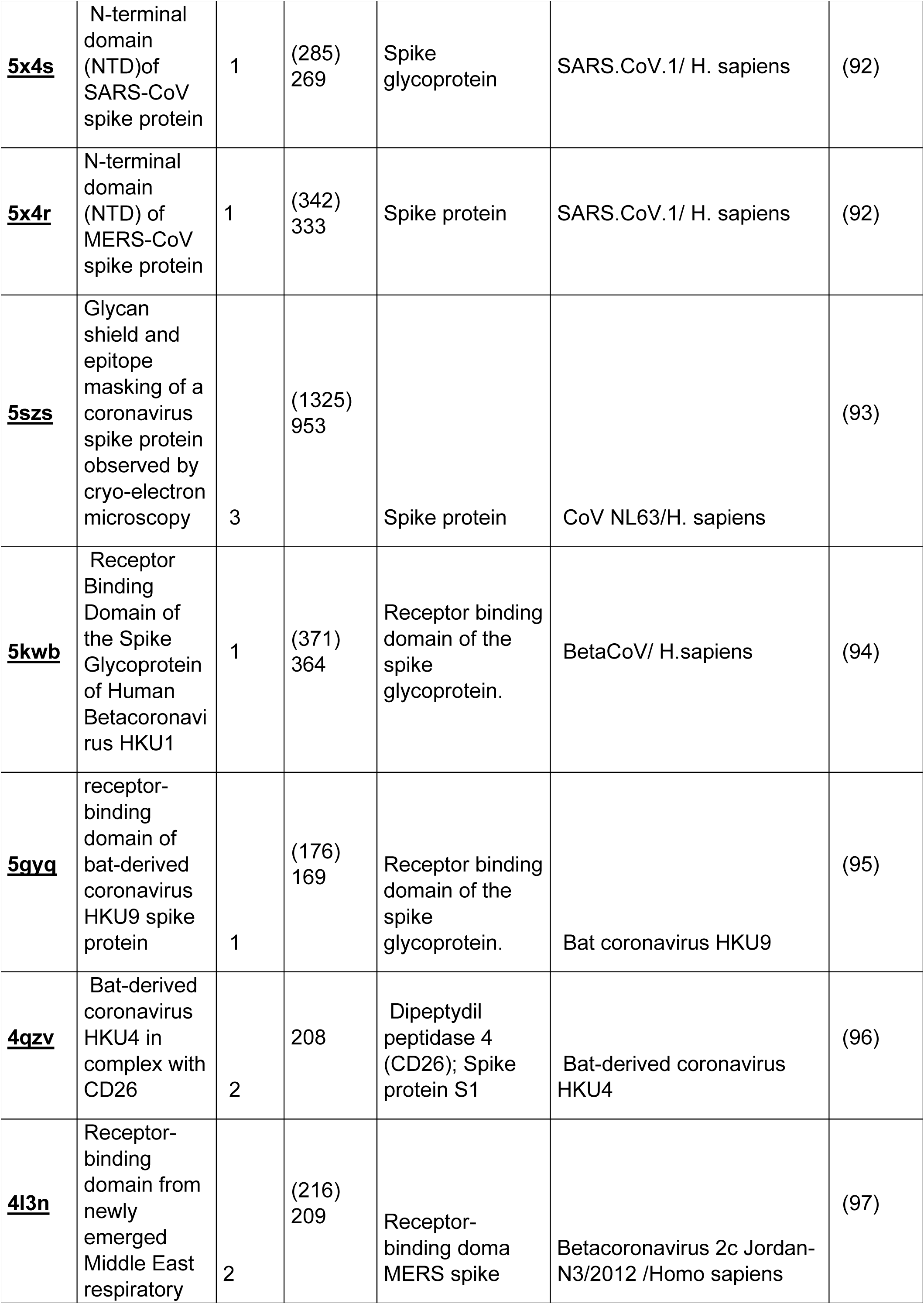
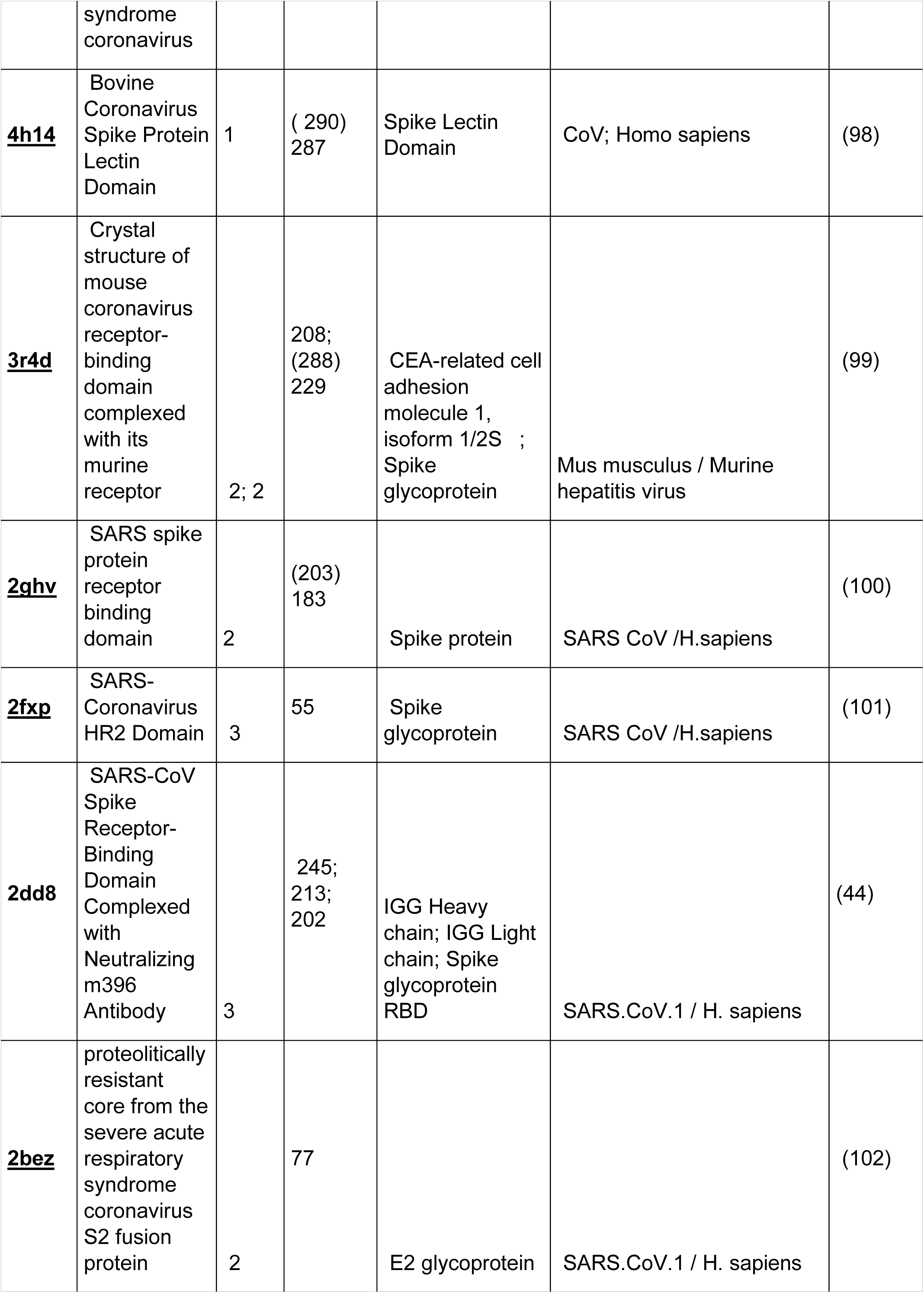
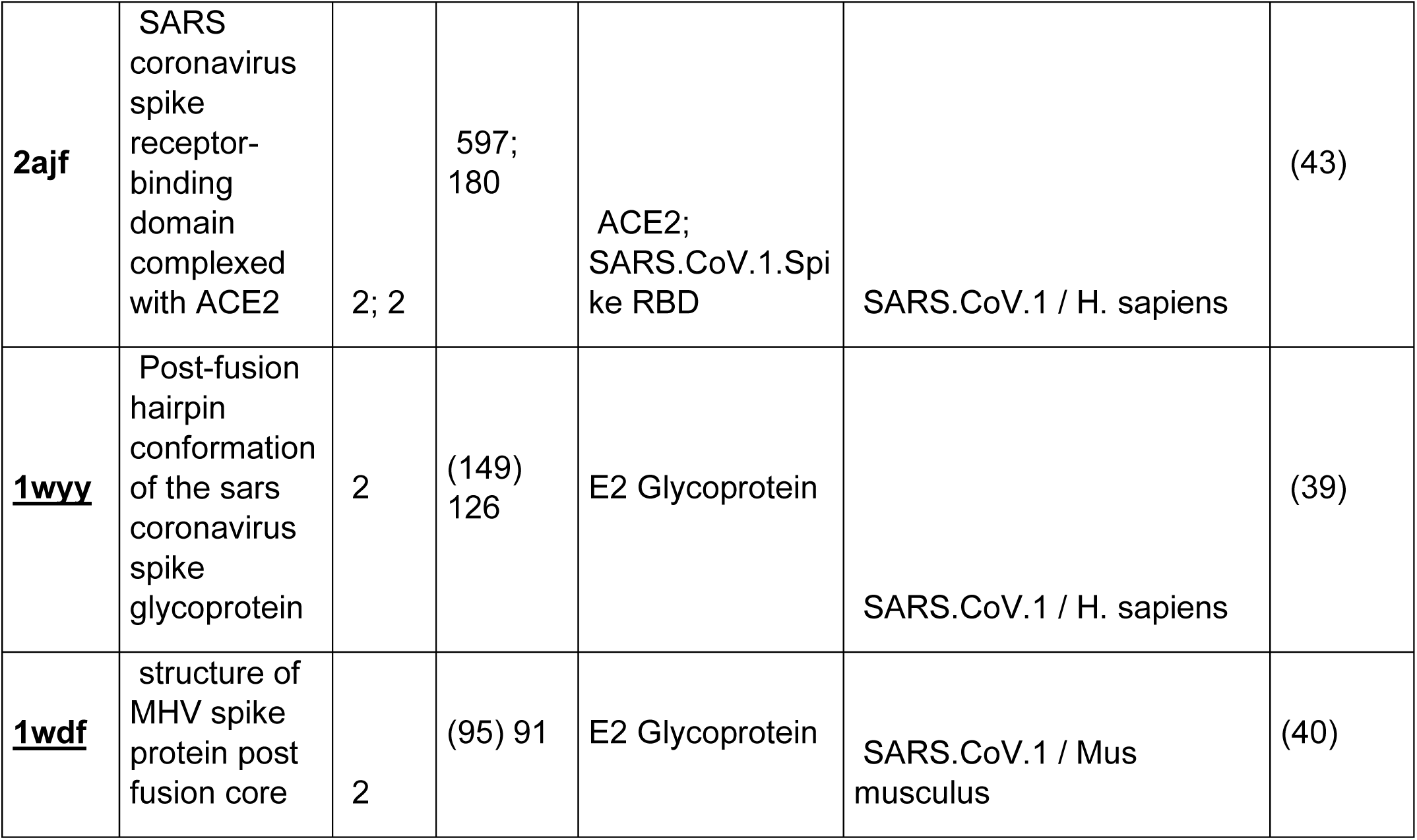
List of the sampled homologous crystallized structures and specific structural features. The listed proteins were sampled by using the folding recognition tools available on pGenTHREADER (http://bioinf.cs.ucl.ac.uk/psipred/) and i-Tasser (https://zhanglab.ccmb.med.umich.edu/I-TASSER/) webservices. SARS-CoV-2 structure with the cited PDB_IDs are also available through the COVID-19/SARS-CoV-2/Resources available on the PDB at the link: https://www.rcsb.org/news?year=2020&article=5e74d55d2d410731e9944f52.

**Supp. Fig. 1.**
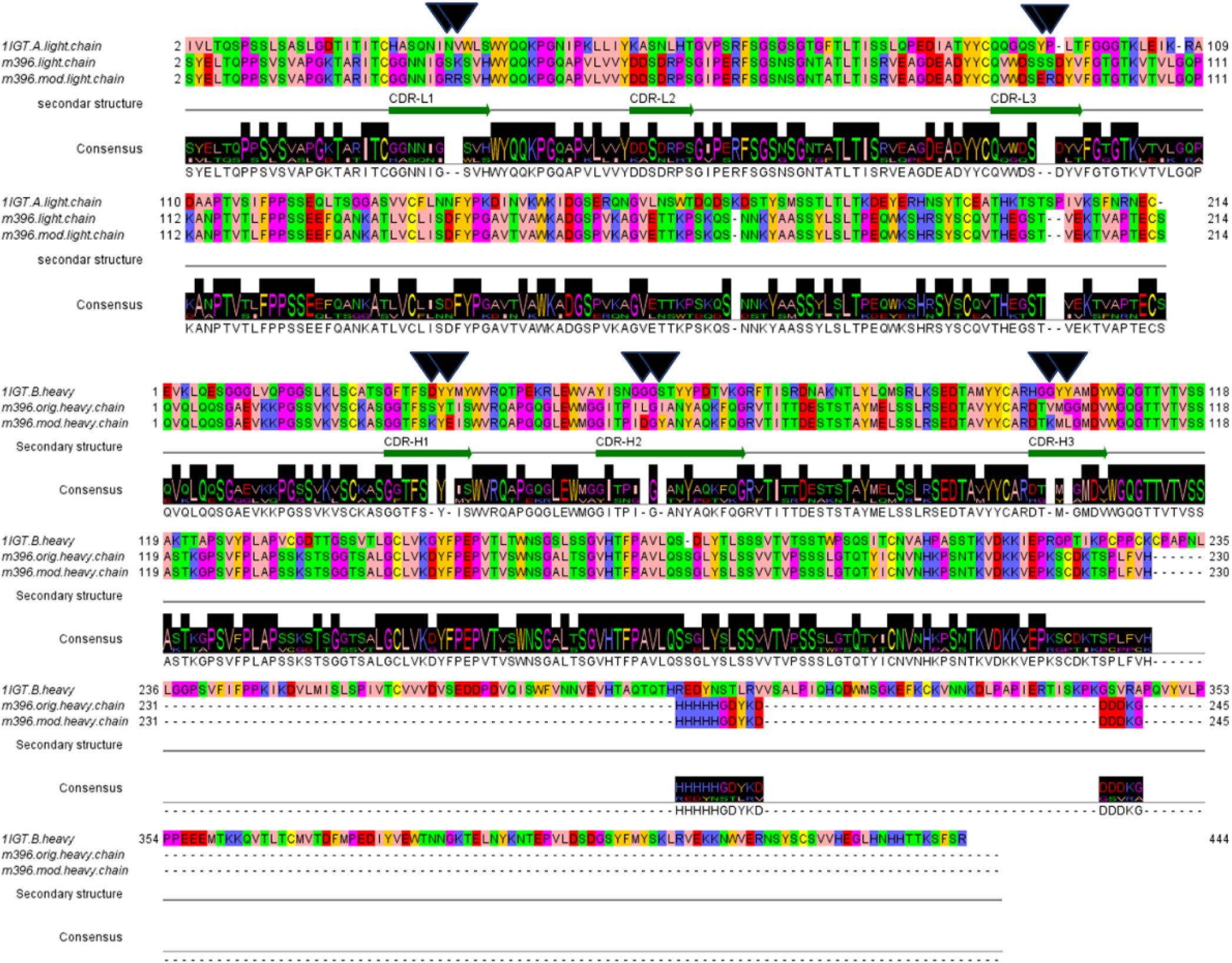
MSA of antigen-binding fragment (Fab) of m396 ab (native and modified) with 1IGT.pdb sequece. (A) Pairwise alignment of the light chains from 1IGT (murine IgG) and m396 antibody (native and modified). (B) Pairwise alignment of the heavy chains of the Fab portions from 1IGT and m396 antibody (native and modified.

**Supp. Fig. 2.**
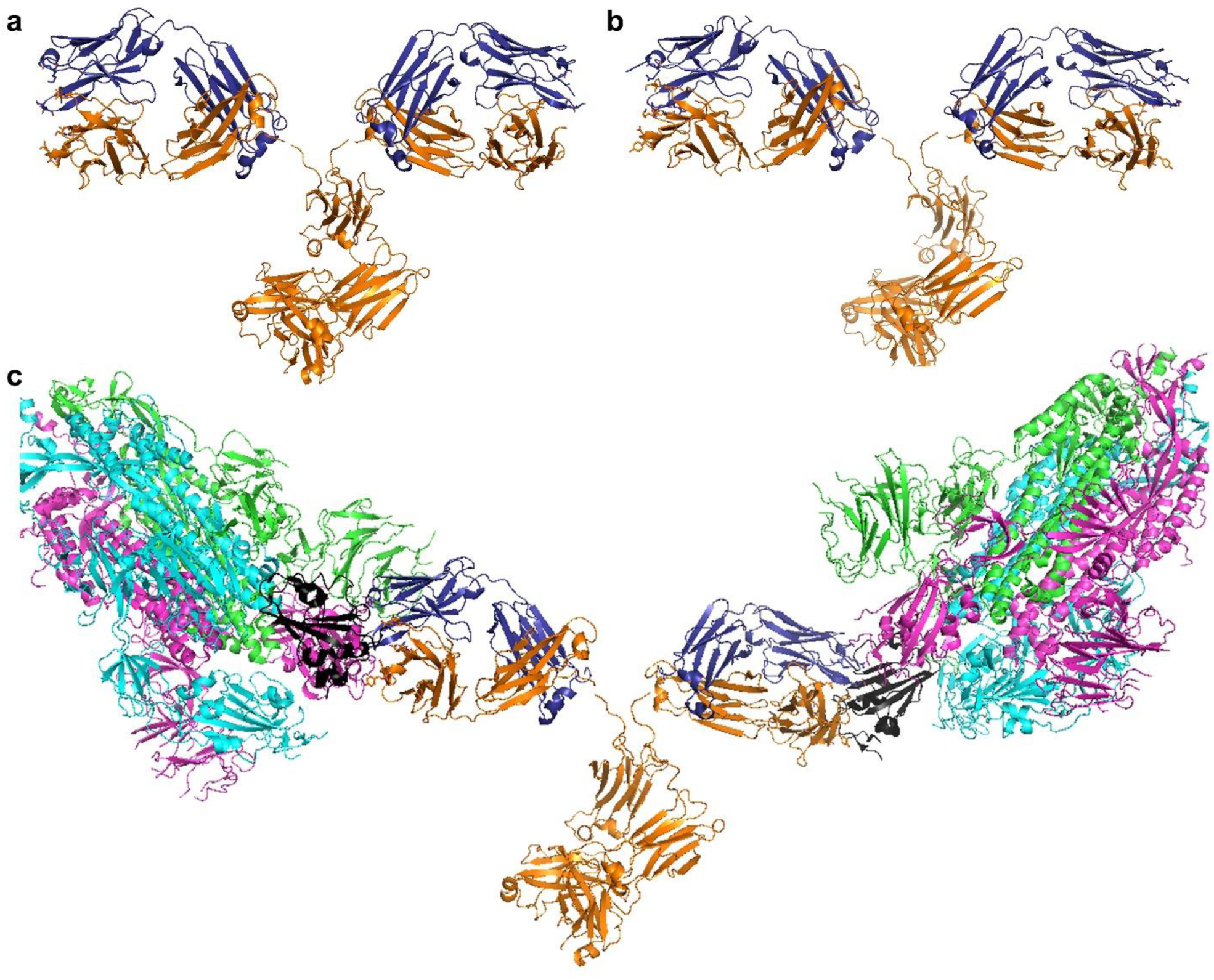
The overall structure of 1IGT antibody used as a protein template for building our chimeric mAb models based on m396 ab. The heavy chains are reported in orange cartoons, and the light chains are reported in blue cartoons, for the native m396 (panel a) and for the modified m396 (panel b). SARS-CoV-2 spike proteins are reported in cyan/magenta/green cartoon representation with the exclusion of RBD reported in black cartoon representation. Breaks in the tertiary structures of the antibody backbones (portion in orange cartoon representation), at the interface of the FAB/Fc portion, indicate sites hosting residues that were ligated after superimposition of the different portions (1igt.pdb and native/mutated m396) for obtaining the final complete 3D model of the proposed antibodies.

